# Variant mutation in SARS-CoV-2 nucleocapsid enhances viral infection via altered genomic encapsidation

**DOI:** 10.1101/2024.03.08.584120

**Authors:** Hannah C. Kubinski, Hannah W. Despres, Bryan A. Johnson, Madaline M. Schmidt, Sara A. Jaffrani, Margaret G. Mills, Kumari Lokugamage, Caroline M. Dumas, David J. Shirley, Leah K. Estes, Andrew Pekosz, Jessica W. Crothers, Pavitra Roychoudhury, Alexander L. Greninger, Keith R. Jerome, Bruno Martorelli Di Genova, David H. Walker, Bryan A. Ballif, Mark S. Ladinsky, Pamela J. Bjorkman, Vineet D. Menachery, Emily A. Bruce

## Abstract

The evolution of SARS-CoV-2 variants and their respective phenotypes represents an important set of tools to understand basic coronavirus biology as well as the public health implications of individual mutations in variants of concern. While mutations outside of Spike are not well studied, the entire viral genome is undergoing evolutionary selection, particularly the central disordered linker region of the nucleocapsid (N) protein. Here, we identify a mutation (G215C), characteristic of the Delta variant, that introduces a novel cysteine into this linker domain, which results in the formation of a disulfide bond and a stable N-N dimer. Using reverse genetics, we determined that this cysteine residue is necessary and sufficient for stable dimer formation in a WA1 SARS-CoV-2 background, where it results in significantly increased viral growth both *in vitro* and *in vivo*. Finally, we demonstrate that the N:G215C virus packages more nucleocapsid per virion and that individual virions are larger, with elongated morphologies.

## INTRODUCTION

The coronavirus disease of 2019 (COVID-19) pandemic originated from the emergence of Severe Acute Respiratory Syndrome Coronavirus 2 (SARS-CoV-2) in late 2019^1^. Subsequent worldwide spread and sustained transmission over the past four years, combined with near real-time access to viral genomic surveillance data, has revealed a detailed picture of SARS-CoV-2 evolution in the human population. Individual mutations that conferred an evolutionary advantage were quickly selected and maintained in viral lineages, leading to the emergence of novel Variants of Concern distinguished by increased immune evasion, transmission, disease burden or infectivity. Understanding the biological function of the individual mutations that led to the emergence of novel variants of concern has important implications for public health consequences as well as our fundamental understanding of basic coronavirus biology.

Most studies concerning these mutations have focused on genetic changes within the spike (S) protein, due to concerns that sufficiently novel spike proteins can allow viral escape from the immune memory induced by vaccination or prior infection^2–5^. However, mutations elsewhere in the viral genome can play key roles in viral replication and pathogenesis^6–8^. The nucleocapsid (N) protein in particular is a genetic ‘hotspot’ for mutations across variants, with several mutations within the flexible linker region associated with increased replication and pathogenesis^8^. Given the many attributed roles of N, including RNA encapsidation^9^, production of viral RNA^10–13^, and viral assembly through interactions with M^14–16^, mutations within this protein are poised to have large impacts on the viral lifecycle. Additionally, N is a highly conserved protein among coronaviruses, making the N protein a versatile diagnostic marker and potential vaccine target candidate^17^.

We have identified a novel cysteine within the linker region of N that was selected and maintained within the SARS-CoV-2 Delta variant. This introduction of a cysteine residue within the nucleocapsid is unique amongst pandemic-causing betacoronaviruses and has major structural implications as it results in the production of a stable N-N dimer linked by a disulfide bond. Here we describe the impact of this novel cysteine within the N protein and the role it plays in viral replication and particle formation. We demonstrate that this cysteine promoted N oligomerization, and that the Delta mutation (G215C) resulted in substantially increased viral replication kinetics in primary differentiated human bronchial cells. The N:G215C mutation also increased viral replication in the nasal washes and lungs of infected Syrian golden hamsters, while paradoxically delaying weight loss. Finally, the N:G215C virus packaged substantially more N per virion and many of the virions displayed elongated morphology. Together, our data suggest that the Delta N:G215C mutation increases levels of nucleocapsid oligomerization which drives increased packaging of N into mature virions and results in significant increases in viral replication both *in vitro* and *in vivo*.

## RESULTS

### Introduction of a novel cysteine in the nucleocapsid linker region of Variants of Concern

The unprecedented access to sequences of SARS-CoV-2 genomes acquired from individual infected people in near real-time throughout the COVID-19 pandemic has revealed a detailed picture of the evolution of this viral pandemic over the last four years. Mutations introduced by the viral RNA-dependent RNA-polymerase led to the emergence of distinct lineages, characterized by suites of different mutations. Variants that met certain public health benchmarks, termed Variants of Concern (VOCs), contained individual mutations in viral proteins that conferred distinct evolutionary advantages. These properties included the ability to evade prior immunity, increased transmission and altered virulence. The Delta variant, which emerged in mid-2021 possessed all the properties above, and contained a series of novel mutations, including key changes in the spike (S) protein^7^. While the function of the mutations in spike has been intensively studied ^4,18,19^, the majority of Delta sub-lineages also contained a unique mutation in the nucleocapsid (N) protein which converted a glycine into a cysteine at position 215 (G215C). This point-mutation dominated the Delta lineages (Fig. 1A) and bioinformatic analysis suggested that its evolutionary advantage was distinct from co-occurring mutations in spike or nucleocapsid^20^.

**Figure 1:**
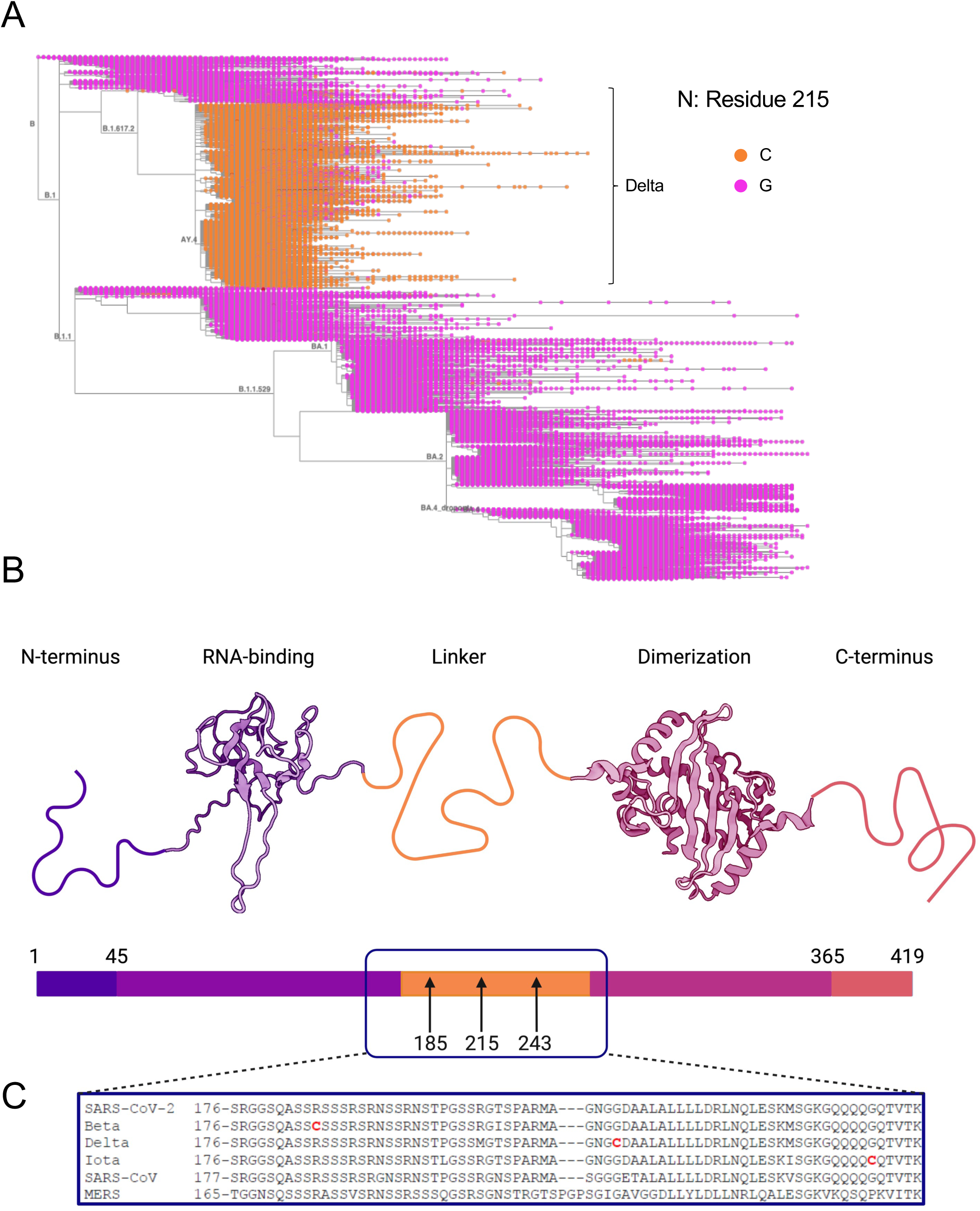
Introduction of novel cysteine residues into the SARS-Co-2 nucleocapsid linker. A) Amino acid identity of position 215 in the SARS-CoV-2 nucleocapsid protein in genomic surveillance data from 7,342,041 sequences in The International Nucleotide Sequence Database Collaboration is shown (Glycine in pink, Cysteine in orange). Sequences were visualized by Taxonium11 on January 10^th^, 2024, at position 215 of the N protein. The tree was rooted to Wuhan/Hu-1 (GenBank MN908947.3, RefSeq NC_045512.2), sequences were added to the tree through the use of UShER12 and Pangolin lineage identity is labelled (B.1617.2 branch shows Delta sequences and B.1.1.529 shows Omicron sequences). B) The SARS-CoV-2 N protein includes an unstructured N-terminus (residues 1-45; in indigo), an N-terminal RNA binding domain (NTD; residues 45-176; in violet, PDB code 6YI3^62^, an unstructured linker (LKR) region (176-263, shown in yellow) which binds to Nsp3, a C-terminal dimerization domain (CTD; residues 263-365; in pink, PDB code: 6WZO^23^, followed by an unstructured C-terminal region (residues 265-419; in peach). C) The sequences of the Nucleocapsid linker region from SARS-CoV-1 (WA1), the Beta, Delta and Iota isolates used in this paper, as well as SARS-CoV and MERS were aligned using Clustal Omega. Cysteines are highlighted in red, and arrows highlight their position in the schematic. Figure created with Biorender.

The G215C mutation sits in the disordered linker region of the nucleocapsid protein, which lies between a N terminal RNA-binding domain (RBD) and a C terminal dimerization domain (Fig 1B). The introduction of a cysteine in the SARS-CoV-2 nucleocapsid is unique amongst zoonotic Betacoronavirus, as neither SARS-CoV, MERS or other SARS-CoV-2 variants’ nucleocapsids contained any cysteines (Fig 1C). Furthermore, while other coronavirus nucleocapsid proteins do contain cysteines, they are largely absent from the linker region (Fig S1). Surprisingly, when analyzing the sequences of our panel of VOCs obtained from clinical specimens, we discovered that two other variant (Beta & Iota) isolates also contained a cysteine within the N protein (Fig 1C). Interestingly, all three of these mutations sit within the intrinsically disordered linker region of N (also termed N3/sN3) between N2 (RNA binding domain) and N4 (dimerization domain). Both our Beta (B.1.351) and Iota (B.1.526) stocks contained novel cysteines in the linker region, R185C (99.7% of reads) and G234C (100% of reads), respectively (Fig 1C). Since the introduction of a cysteine residue would allow for the formation of a new disulfide-bonded N-N dimer complex, we predicted that this mutation could have major impacts on the secondary, tertiary and/or quaternary protein structure of the nucleocapsid protein.

### Cysteine in the N linker promotes the formation of a stable N-N dimer

To test if these novel cysteines would make a more stable N-N dimer, we visualized the N from our panel of variant isolates by western blot under non-reducing conditions. We infected VeroE6-TMPRSS2 cells with wildtype virus (ancestral SARS-CoV-2 from the WA1 infectious clone) as well as low passage stocks of the Alpha, Beta, Gamma, Delta, Epsilon, Iota, Mu, and Omicron variants isolated from clinical samples. All viruses produced a band of the expected molecular weight (∼47 kDa) for the N monomer, and the majority also showed a series of truncation products (indicated with <) that we hypothesize to be caspase cleavage products^21^ (Fig 2A). As predicted, the three variants which contained the novel cysteine residue (Beta, Delta, Iota) produced a second band detected at twice the molecular weight of the expected N monomer (Fig 2A, see *). Of the three variants, Delta (G215C) produced the greatest level of dimerized-N, with Iota (G243C) and Beta (R185C) each producing slightly lesser amounts (Fig 2B). While it is known that coronavirus N proteins typically form dimers via their dimerization domains, this is mediated by non-covalent bonds and the canonical N-dimer was not seen by SDS-PAGE gel/western blot for viruses that lacked cysteines (WT, Alpha, Gamma, Epsilon, Mu and Omicron).

**Figure 2:**
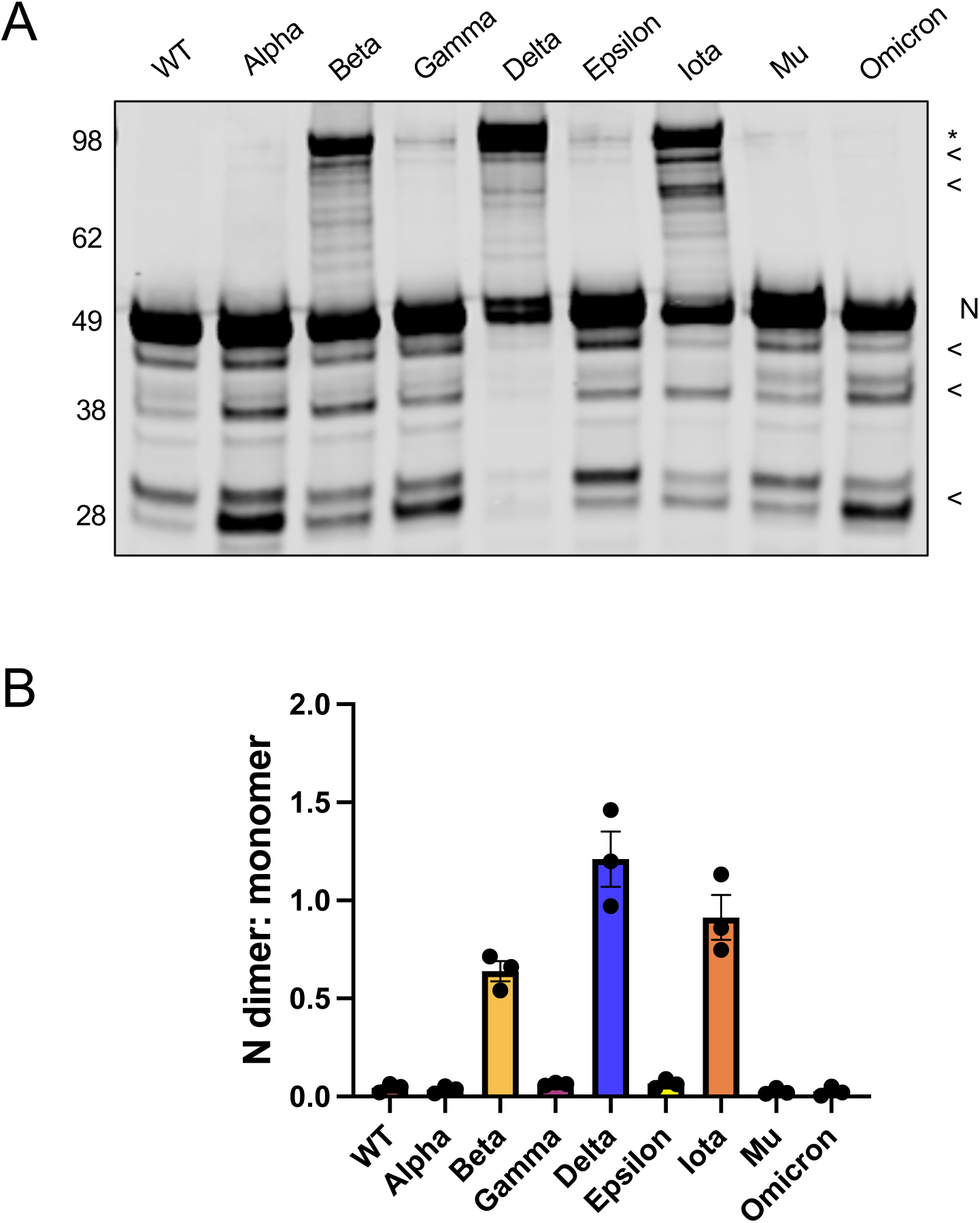
A disulfide-bonded nucleocapsid dimer is formed in several Variants of Concern. VeroE6-TMPRSS2 cells were infected with the indicated SARS-CoV-2 variants (or WA1, termed Wildtype) at an MOI of 0.01 for 24 hours. Unreduced cell lysates were visualized by SDS-PAGE and western blot using antibodies recognizing SARS-CoV-2 N. The presence of a ∼100 kDa band recognized by the SARS-CoV-2 N antibody in the Beta, Delta and Iota variants is indicated by a *. (A) A representative gel and B) the relative levels of N dimer to monomer from 3 independent biological replicates is shown. Mean +/- SEM is plotted, individual data points for each experiment are superimposed onto bar graphs for each condition.

To further test whether this higher molecular weight band represented a disulfide-bonded form of N-N dimer we performed several additional experiments. First, we prepared samples under strong reducing conditions (10 mM DTT), where we observed the higher molecular weight band was eliminated (Fig S2A). To ensure that this disulfide bond truly occurred within infected cells and was not a post-lysis artifact, lysates were harvested in the presence of n-ethylmaleimide (NEM). NEM binds irreversibly to free cysteines and prevents the formation of post-lysis disulfide bonds, and the band belonging to the putative N dimer remained visible in these conditions (Fig S2B). To confirm the putative dimer was not an artifact of the monoclonal antibody used, we probed lysates with a second independent antibody (Fig S2C/D). Slight differences in cleavage products were seen with the two antibodies (see *), confirming that the antibodies recognized different epitopes, but the higher molecular weight band was visible in both conditions (Fig S2C-E). To ensure that the higher molecular band did not represent a disulfide bond formed between N and a cellular or viral protein of equivalent weight, we immunoprecipitated N from cells infected with WT or Delta SARS-CoV-2 and performed mass spectrometry on tryptic peptides from gel slices cut from the regions corresponding to the monomer and putative ‘dimer’ (Fig S2F). In the Delta infected sample, most peptides detected in the ‘dimer’ gel slice corresponded to N, and roughly half the total N peptides were detected in the ‘dimer’ vs ‘monomer’ slice (Fig S2G). Furthermore, there were no cellular peptides of similar abundance found in the ‘dimer’ slice, with the most abundant cellular protein found at >1/5^th^ the levels of N (Table S1). Given the structural location of these cysteines in the linker region, and the fact that N is known to oligomerize into higher order structures^22,23^, we think it is likely that the introduction of this disulfide bond promotes the formation of a tetramer or higher order structure by stabilizing the bond between two non-covalently-bonded dimers.

### N dimer stability dependent on novel cysteines in linker region

As the production of a disulfide bond in the reducing environment of the cytoplasm is unusual, we next determined whether the formation of this stable N-N dimer required the context of infection. SARS-CoV-2 replication produces many membranous compartments with limited cytoplasmic access that could shield N during authentic infection. We created plasmid expression constructs of the WT, Delta, Beta and Iota nucleocapsid sequences and transfected these into HEK-293T cells, in conjunction with constructs where each cysteine was mutated back to the canonical residue in the WT sequence (Delta C215G, Beta C185R, Iota C243G). While all three variant constructs (Delta, Beta, Iota) were capable of producing stably dimerized N (Fig 3A), Delta produced this product to the highest levels while the Beta construct did not consistently form visible, stable dimer (Fig 3B). In each case, when the cysteine was reverted to its original amino acid the stabilized dimer was not made (Fig 3A/B). These data suggest that, at least in the case of Delta and Iota, the presence of other viral proteins/RNA and the formation of double membrane vesicles (DMVs) are not required for stably dimerized N formation, and the presence of a cysteine at 215/243 in the Delta/Iota backgrounds is sufficient.

**Figure 3:**
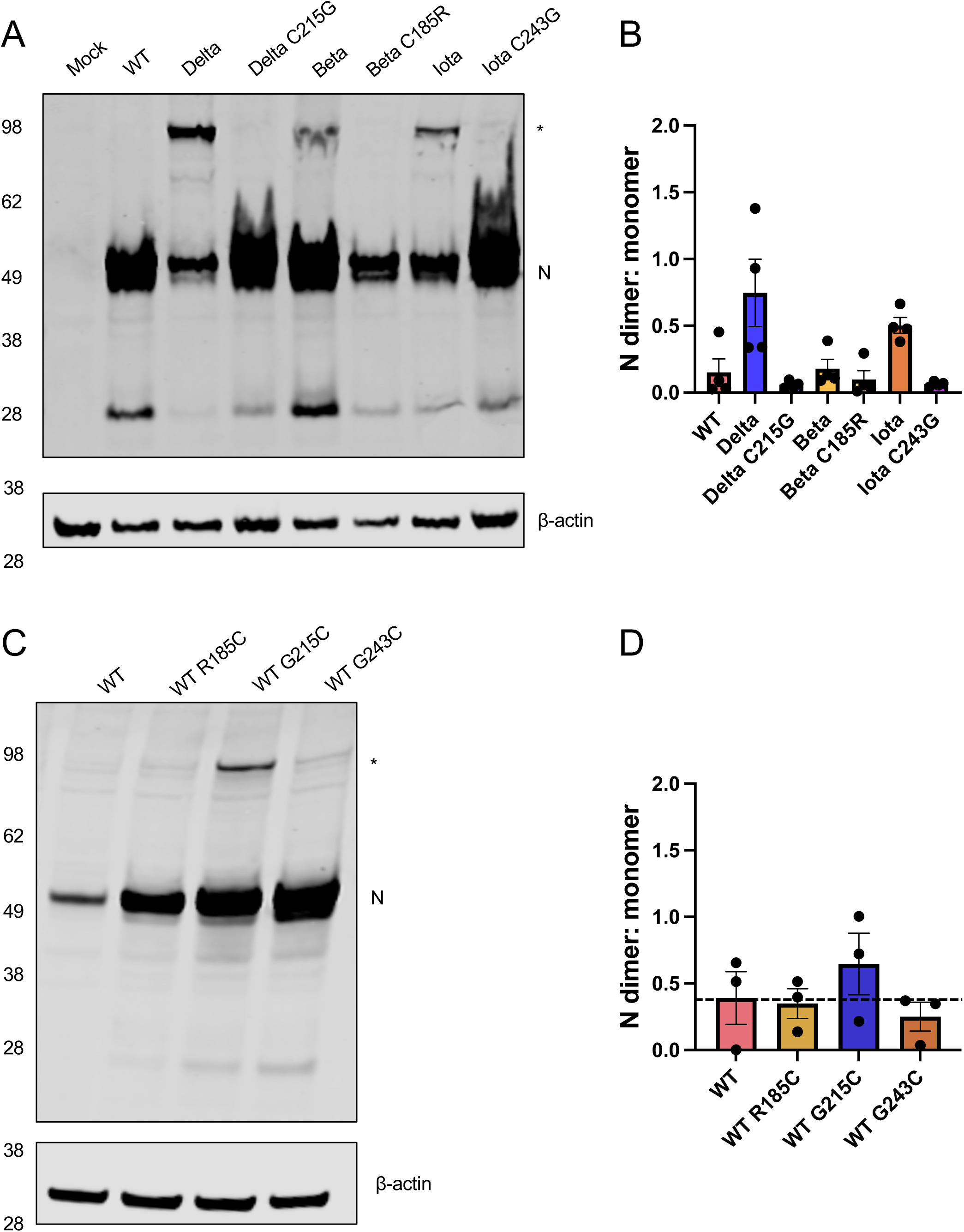
N-linker cysteine residues are necessary and sufficient for stable dimer formation. (A) HEK-293T cells were transfected with plasmids encoding the N protein of the indicated SARS-CoV-2 variant. In addition, cells were transfected with constructs in which each cysteine was changed back to the parental residue in the WT (WA1) sequence. 24 hours post transfection, unreduced cell lysates were visualized by SDS-PAGE and western blot using antibodies recognizing SARS-CoV-2 N and β-actin. (B). The ratio of N monomer to dimer bands seen in the western blot shown in A) was quantified. C) HEK-293T cells were transfected with plasmids encoding the WT N protein or constructs introducing a cysteine at positions 185, 215 or 243 and (D) the ratio of monomer to dimer bands was quantified. The presence of a ∼100 kDa band recognized by the SARS-CoV-2 N antibody corresponding to the stable dimer is indicated by a *. A representative western from four (A/B) or three (C/D) independent biological experiments is shown, probed for N (red) and beta-actin (green). Mean +/- SEM is plotted, individual data points for each experiment are superimposed onto bar graphs for each condition.

### Impact of the N:G215C mutation on viral growth *in vitro* and *in vivo*

As bioinformatic evidence suggests that Delta variants containing the G215C mutation outcompeted those that contained the ancestral glycine at that position^20^ we wanted to examine the impact of the G215C mutation on viral growth kinetics. Since WT SARS-CoV-2 (WA1/2020) does not produce the stably dimerized N in infection (Fig 2A), we investigated if the introduction of a cysteine at 185, 215 or 243 (Beta, Delta, Iota) was sufficient to mediate stable dimer formation in a WT background. Using plasmid constructs, we transfected HEK-293T cells with either WT N, or WT N containing a single point mutation (R185C, G215C, and G234C). When visualized via western blot in non-reducing conditions, only the construct which contained the Delta G215C mutation could produce the stabilized dimer, suggesting that this mutation is both necessary and sufficient for stable dimer formation (Fig 3C,D).

As inserting the G215C mutation in the WT background was sufficient to confer dimer formation, we used a well-established reverse genetics system^24^ to rescue an infectious clone containing the G215C nucleocapsid mutation (N: G215C) in the SARS-CoV-2 WA1 backbone (Fig 4A). We used a neon-green reporter virus in the WA1 background (mNG SARS-CoV-2), which is genetically identical to WA1 except that ORF7 has been replaced with a neon green fluorescent reporter). Notably, mNG SARS-CoV-2 is attenuated compared to the parental WA1, due to the replacement of ORF7a with the mNG reporter^8^. As expected, the N:G215C virus produced the stable N dimer in infected Vero-TMPRSS2 cells under non-reducing conditions (Fig 4B). In a multi-cycle growth curve, the WT and N:G215C viruses grew to identical peak titers, with very similar growth kinetics. We did note that at the earliest timepoint (7hpi), the WT virus shows an ∼1 log drop in viral titer (representing the loss in infectivity of the original inoculum), while the N: G215C virus did not display such a drop (Fig 4C), though the biological significance (if any) of this observation is unclear.

**Figure 4:**
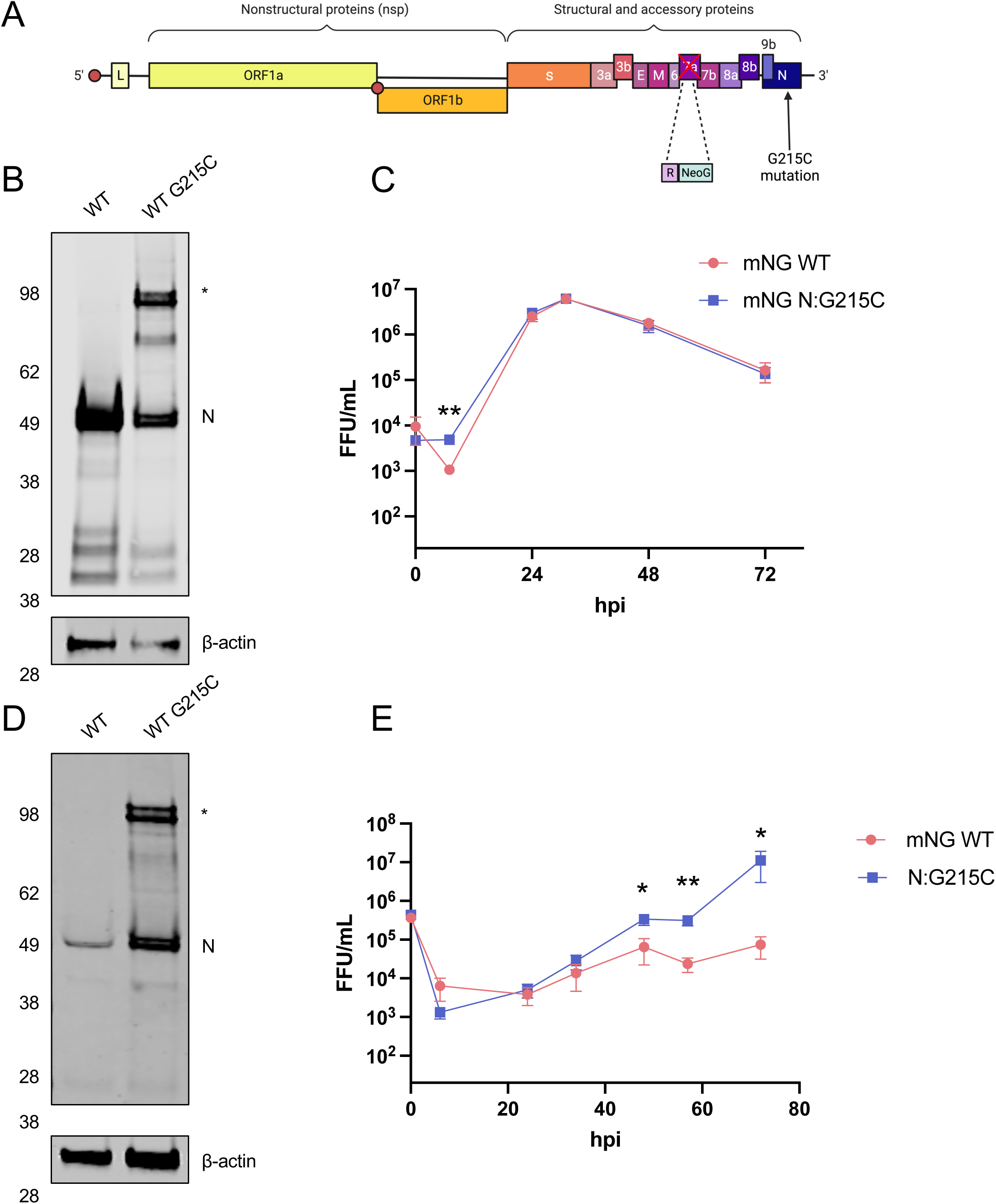
The introduction of N:G215C in a WA1 infectious clone recapitulates stable N dimer formation and displays altered growth kinetics in HBECs. A) A schematic of the SARS-CoV-2 genome encoding the mNG infectious clone system is shown, including the replacement of ORF7a with mNeon Green, and the introduction of the G215C mutation in N. B) VeroE6-TMPRSS2 cells were infected, or mock infected, with the WT or N:G215C viruses at an MOI of 0.0005 for 1 hour. 72 hpi, unreduced cell lysates were collected and visualized by SDS-PAGE and western blot using a rabbit anti-SARS-CoV-2 N and beta-actin. The presence of the higher MW (∼100 kDa) band seen in the N:G215C virus is indicated with a *. A representative gel from three independent biological experiments is shown. (C) In addition, viral supernatants were collected at 8, 24, 32, 48, and 72 hours post infection and titered by focus forming assay (FFU; focus forming unit). D) Human bronchial epithelial cells were infected with WT or N:G215C viruses at an MOI of 0.5 for 1 hour. D) Unreduced cell lysates were collected and processed for western blot as in (B), while E) viral supernatants were collected sequentially from the same well at 8, 24, 32, 48, and 72 hours post infection and titered by focus forming assay (FFU; focus forming unit). Mean +/- SEM is plotted [* (p<0.05), ** (p<0.01)], N=6 from three biological experiments (C), N=12 from four biological experiments (E) is shown. Limit of detection (LoD) is 20 FFU/ml and the Y-axis minimum is set as the LoD.

Next, as VeroE6 cells are not a physiologically relevant target for SARS-CoV-2 infection, we examined viral growth kinetics in primary differentiated human bronchial epithelial cells (HBECs), grown in transwells on an air-liquid interface (ALI). As expected, the N:G215C virus produced the stable N dimer in infected HBECs when harvested under non-reducing conditions (Fig 4D). Notably, in multi-cycle growth curves the mNG N:G215C virus had improved growth kinetics, with a peak titer more than 100 times greater than the mNG WT virus (Fig 4E). These data suggest that the stably dimerized form of N conveys a particular advantage to viral replication in primary differentiated human bronchial cells.

Next, we determined the effect of the N:G215C mutation *in vivo* in the Syrian Golden Hamster model of SARS-CoV-2 infection. Three to four week-old male hamsters were inoculated intranasally with either PBS (mock), 10^4^ PFU of the WT neon green reporter SARS-CoV-2 (mNG WT) or 10^4^ PFU of the neon green reporter SARS-CoV-2 containing the N:G215C mutation (N:G215C). Animals were monitored for weight gain/loss daily for seven days, and cohorts of five animals underwent nasal washing followed by euthanasia to obtain tissues to determine viral loads in the lung at both day 2 and day 4 (Fig. 5A). On day seven, surviving animals were euthanized and lung tissue collected for virological and histopathological analysis. While animals infected with the N:G215C mutation showed significant weight loss, relative to control WT virus, the kinetics were delayed with peak disease achieved at day 5-6 post infection, 1-2 days after WT infection (Fig 5B). Strikingly, despite delayed weight loss, viral replication was increased with the G215C mutation. Modest but significant increases in viral titers were observed in the nasal washes at day 4, and a sustained 10-fold increase over WT titers in the lungs throughout the infection (Fig 5C-D). Together, despite the kinetic delay, the N:G215C mutant caused similar overall weight loss and augmented viral replication.

**Figure 5:**
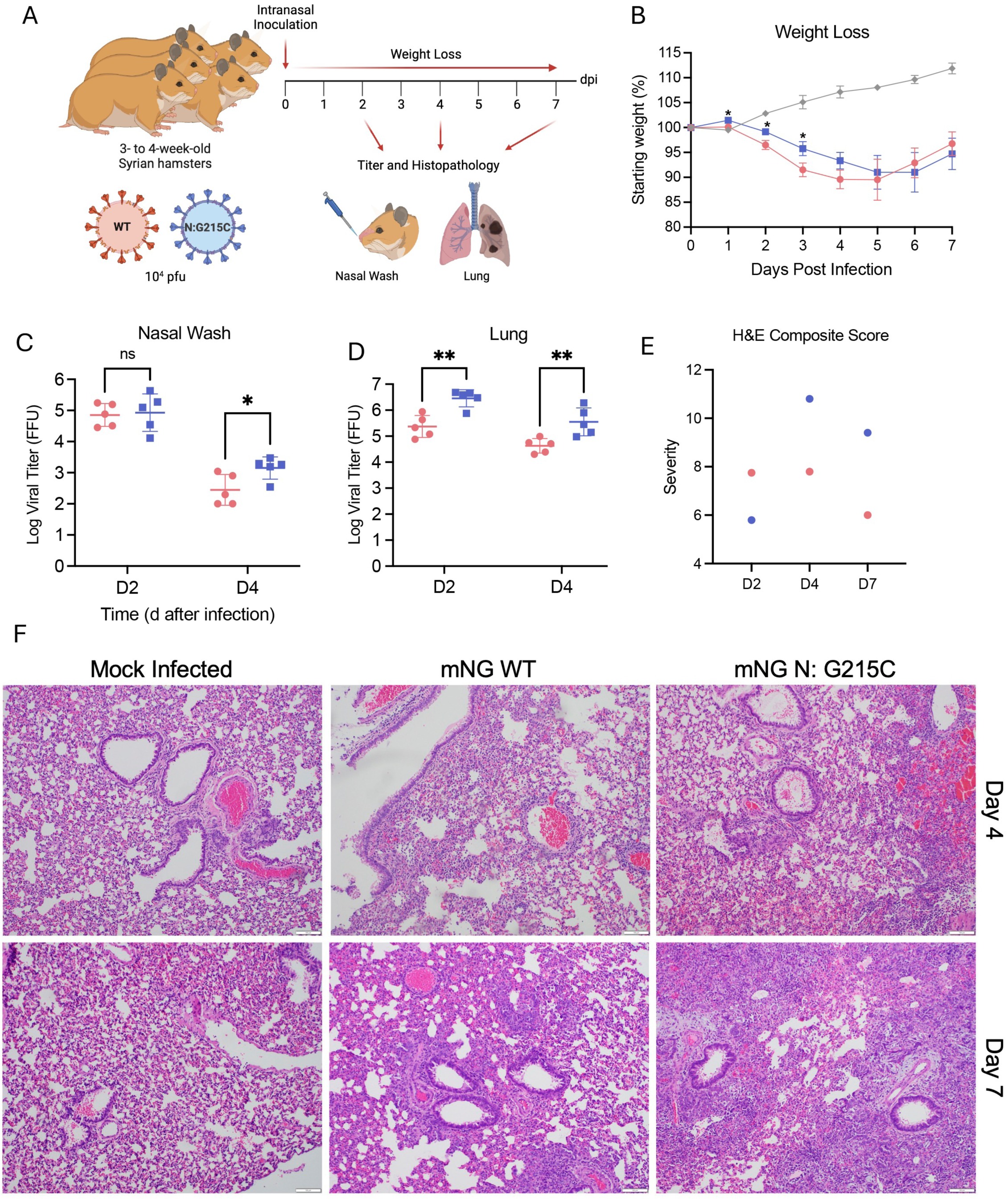
The N:G215C mutation increases viral growth in nasal washes and the lungs of Syrian golden hamsters as well as immune infiltration and damage. A) Three- to four-week old male hamsters were intranasally infected with PBS alone (mock - grey) or 10^4^ FFU of WT (pink) or N:G215C (blue) m Neon Green (NG) SARS-CoV-2. B) Weight-loss of infected animals was monitored daily for 7 days post infection. On days 2 and 4 post infection, titer in the nasal wash (C) and right lung (D) were determined. On days 4 and 7 post infection, left lung tissue was harvested, fixed, cut into 5 µM section, stained with hemoxylin and eosin, scored for pathological severity (E) and imaged (F). For weights, graphs represent mean weight change +/- SEM. For viral titers, lines represent mean viral titer +/- SD. Statistical differences were determined by student’s T-test with * (p<0.05), ** p<0.01). Scale bars are 100 μm.

Examining histopathology, the N:G215C mutant had modest changes in antigen staining, but increased infiltration and damage relative to WT control virus. At day 2, both WT and N:G215C had similar antigen distribution and scores (Fig. S3 A-B). By day 4, N:G215 had a modest increase in overall antigen staining mostly driven by significant differences in airway staining. Viral staining was cleared in both WT and mutant infected animals by day 7 post infection. Examining immune infiltration and damage, lesions were of similar composition and size at day 2 for both groups, but more severe in WT animals (Fig. 5E-F). However, at day 4, N:G215C had increased infiltration and damage compared to WT infected animals. While interstitial pneumonia, bronchiolitis, periarterial edema was common in both groups, N:G215C infected mice were consistently observed to have epithelial cytopathology and subendothelial mononuclear cells. Similarly at day 7, N:G215C maintained evidence of significant damage with continued peribrochiolitis and epithelial cytopathology; in contrast, WT infected hamsters had reduced overall damage scores. Together, the results demonstrate that despite similar weight loss, the N:G215C mutant infected animals have increased viral antigen accumulation and damage in the lung as compared to control.

### Wildtype SARS-CoV-2 containing the G215C mutation packages more nucleocapsid protein per virion and displays oblong morphology

The SARS-CoV-2 nucleocapsid, like nucleocapsids of other Betacoronaviruses, is a highly multifunctional protein. Like other coronavirus nucleocapsids, it is thought to play key roles in the packaging of viral RNA^25^, the production of viral RNA through interactions with the replication-transcription complex^10–13^, and the antagonism of the innate immune response^26–28^. To better understand why the N: G215C mutation was important at a molecular level, we looked at where in the viral life cycle the stably dimerized form of N was observed. We first observed the stable N-dimer in lysates of cells infected with the Beta, Delta and Iota variants (Fig 2A), though cells transfected with the Delta (and to lesser extent Iota) nucleocapsids were able to form the durable N-dimer even in the absence of other viral machinery (Fig 3A, B). As we had observed a gradient of dimer formation depending on where the cysteine mutation was located (215>243>185; Fig 2B, 3B), we next explored whether the stable G215C N-dimer was found at highest levels in transfected cells, infected cells, or in concentrated extracellular virions.

Accordingly, we measured the ratio of N dimer:N monomer visible on a western blot of samples collected under non-reducing conditions from transfected HEK-293T cells, infected Vero-TMPRSS2 cells, or extracellular virus concentrated by polyethylene glycol (PEG) precipitation (Fig 6A-C). Interestingly, we noted that the dimer was enriched in extracellular virus (Fig 6C, D), though infection (vs transfection in isolation) also appeared to promote formation of the stably dimerized N (Fig 6B vs A).

**Figure 6:**
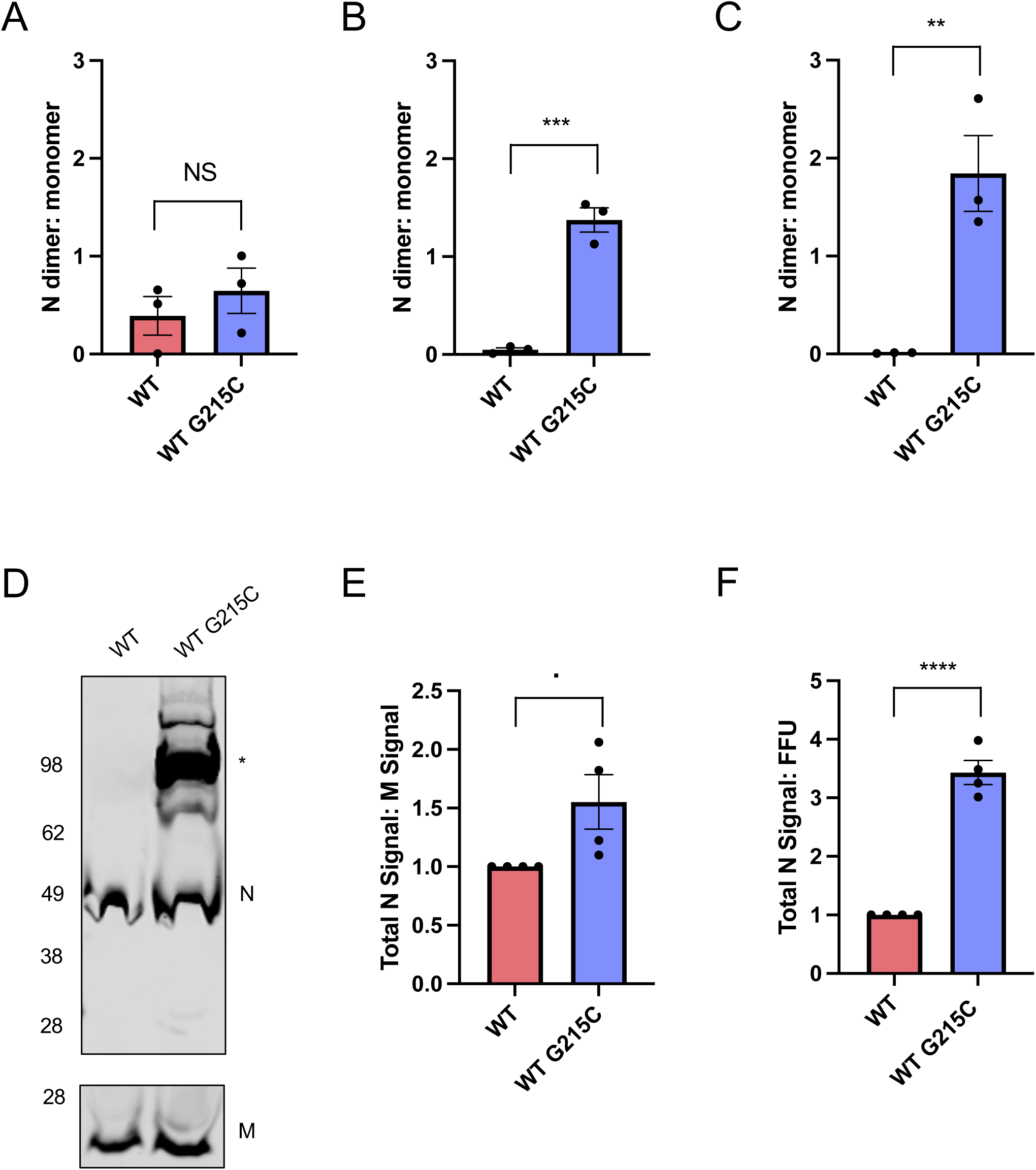
Stably dimerized N is found preferentially in virions and the G215C mutation results in increased viral packaging of N. (A) HEK-293T cells were transfected with plasmids encoding the WT or G215C N protein. B) Vero-TMPRSS2 cells were infected with the WT or N:G215C viruses at an MOI of 0.0005 for 1 hour. C) High titer viral stocks of the WT or N:G215C viruses were concentrated by binding to 10% polyethylene glycol (PEG) then centrifugated at 10,000G for 30 minutes at 4°C. Unreduced lysates were collected 24 hours post transfection, 72 hours post infection or directly from the concentrated viral pellet, visualized by SDS-PAGE and the ratio of N monomer to dimer was quantified. D) A representative western of (C) is shown, probed for N and M. E) The ratio of N to M (normalized to the WT N:M ratio for each replicate) is shown, as well as (F) the ratio of N to FFU for each stock (again normalized to the N:FFU ratio in WT virus for each replicate). N=3 for A, B,C, N=4 for D, E, F. Mean +/- SEM is plotted, individual data points for each experiment are superimposed onto bar graphs for each condition [. (p=.05-.1), * (p<0.05), ** (p<0.01), *** (p<.001) or **** (p<.0001).].

As this enrichment of dimerized N in virions suggested a potential role in encapsidation, we next compared the incorporation of total levels of N to levels of M in the WT vs N: G215C viruses (Fig 6D, E). Finally, we compared the levels of N to the number of infectious focus forming units (FFU) for the two viruses (Fig 6F). In both cases the N: G215C virus appeared to over-incorporate N in virions, compared to another structural protein (M) or infectious units, suggesting that the stably dimerized N has increased encapsidation activity compared to the WT nucleocapsid protein.

In order to examine these apparent changes in encapsidation and packaging on a single virion level, we performed thin section transmission electron microscopy (TEM) on Vero-TMPRSS2 cells that were infected with the WT or N:G215C virus. We examined the morphology of mature virions that had completed budding into intracellular compartments and were no longer attached to the cellular membrane, assessing virion shape as well as the amount and arrangement of internal nucleocapsid structures inside individual virions (Fig 7). The WT virions were largely round and ∼60 μM in diameter, with electron dense complexes likely representing ribonucleoprotein (N+RNA) complexes packed inside (Fig 7A). The N:G215C virus produced some spherical particles similar to WT virions, but it also produced a substantial fraction of virions that displayed oblong or elongated morphologies, and were larger than the WT virions in circumference (Fig 7A,B, Fig S4). These virions also appeared to package more ribonucleoprotein complex structures than the smaller spherical virions, in agreement with our previous biochemical analysis of bulk virions.

**Figure 7:**
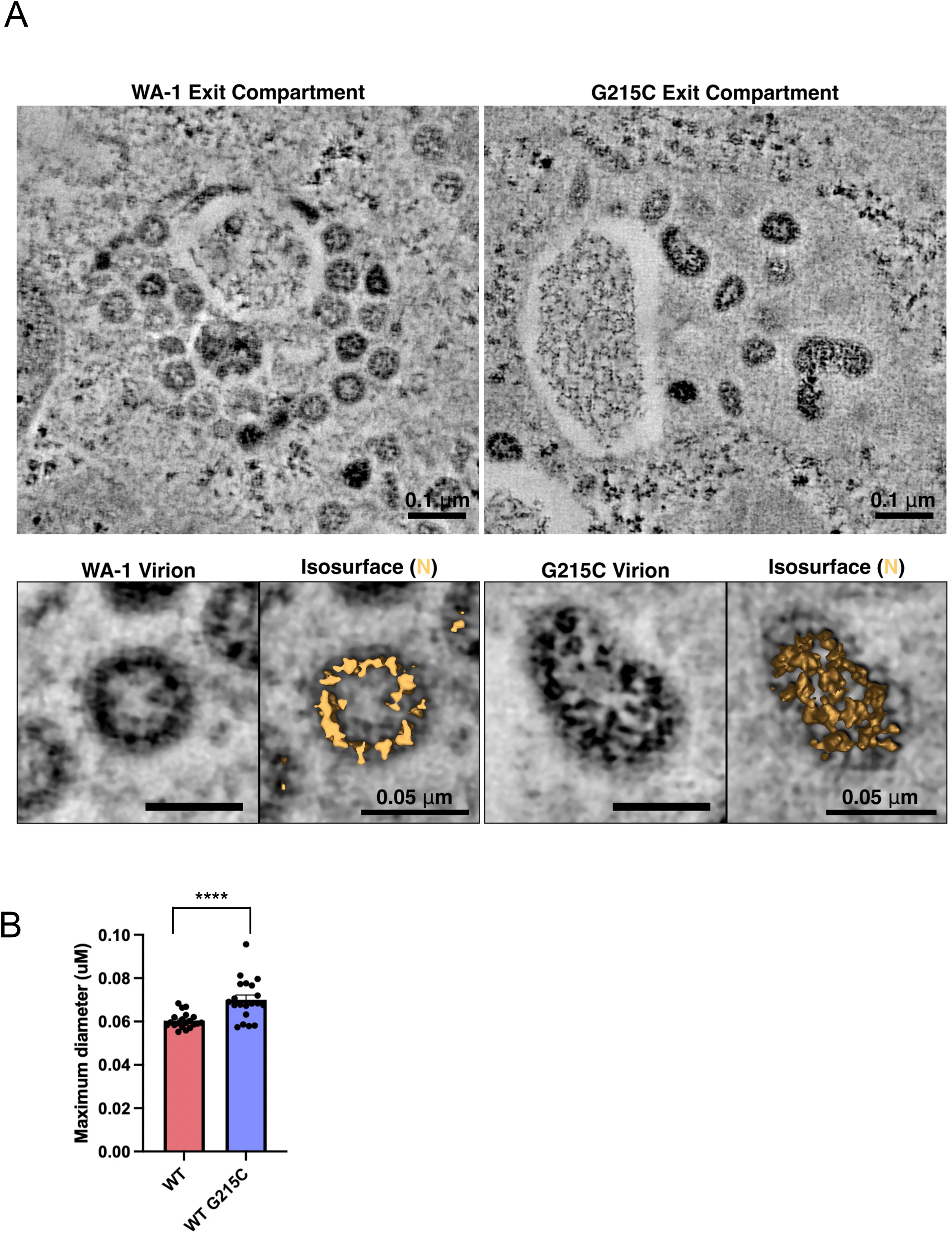
N:G215C virions are enlarged and show over-incorporation of N. A) Vero-TMPRSS2 cells were infected with WT or N:G215C viruses at an MOI of 0.1. The following day cells were prepared for electron microscopy by high-pressure freezing and freeze-substitution, then sectioned and imaged by dual-axis electron tomography. Virus-containing exit compartments were located in both samples, and virions that had completely separated from cellular membranes were selected and analyzed in 3D in order to determine the structure of intact virions and the arrangement of internal ribonucleoprotein complexes. B) The maximum diameter of 20 randomly selected virions for each virus was measured (see Fig S4). Mean +/- SEM is plotted, individual data points for each experiment are superimposed onto bar graphs for each condition, **** (p<.0001).

## DISCUSSION

In this study, we investigate the role of unique cysteine residues that were inserted into the SARS-CoV-2 nucleocapsid linker domain in several Variants of Concern. We demonstrate that these cysteines can form stable disulfide bonds that link nucleocapsid proteins together, outside of the canonical dimerization domain. Sequencing data from public health surveillance efforts suggested that the insertion of a cysteine at position 214 (Lambda) and 215 (Delta) were maintained in these two lineages (Fig 1), and our data support the observation that the G215C mutation is beneficial to the virus *in vitro* (Fig 4E) and *in vivo* (Fig 5B, C). While Betacoronaviruses are known to form dimers, this process is generally thought to occur via a dimerization domain in the C terminus of the protein and importantly is not normally the result of a disulfide bond. Due to the location of the two conserved cysteine residues (214 & 215) in the center of the flexible N linker, and prior data suggesting the linker can play a role in oligomerization^20,23^, we propose that these mutations form disulfide bonds between pairs of N-N dimers and mediate higher order N oligomerization (Fig 8). Increasing the affinity of the inter-linker interactions that mediate oligomerization could have the effect of shifting the balance to N towards higher levels of oligomerization.

**Figure 8:**
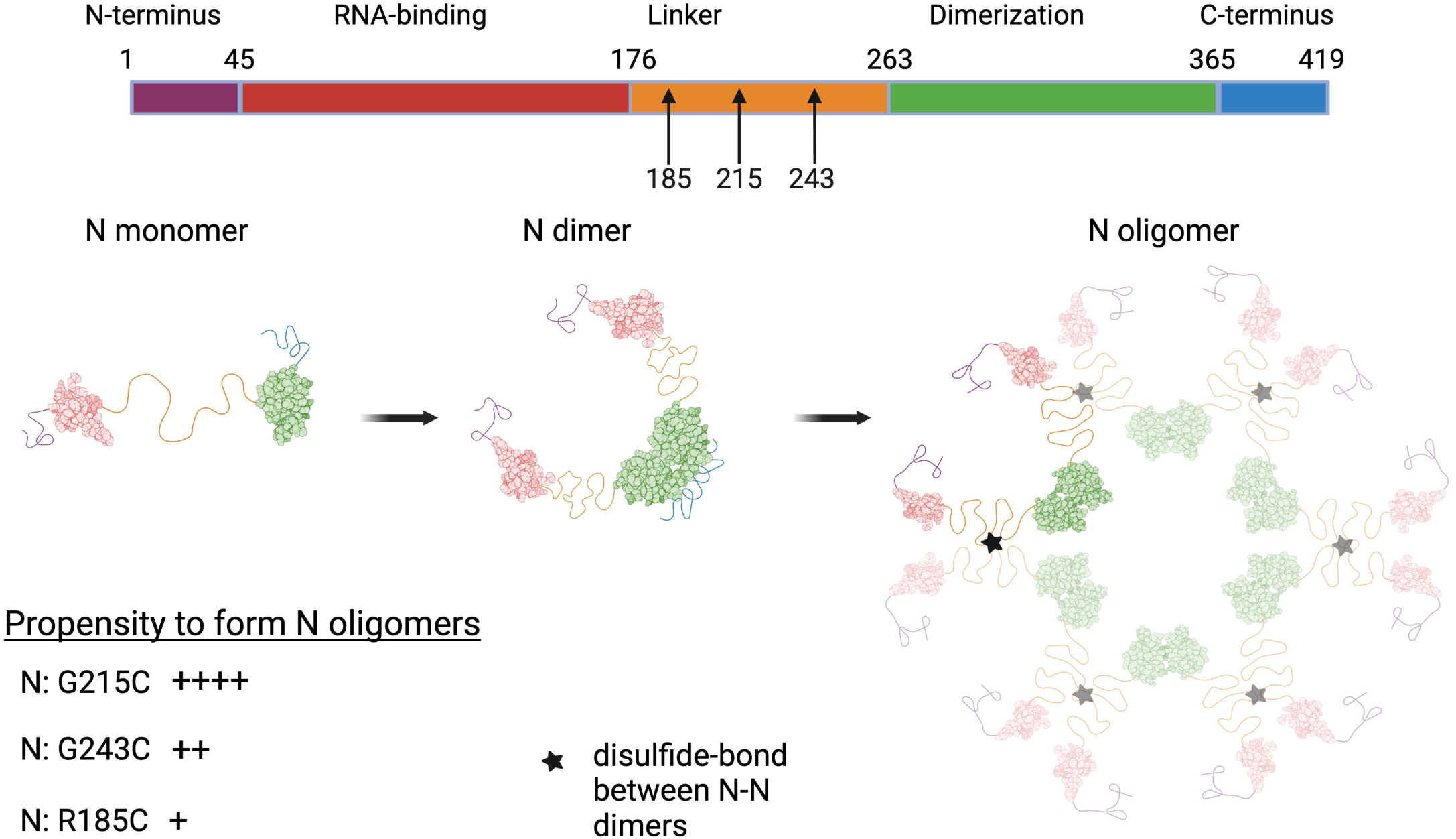
Mutations in the nucleocapsid linker may alter the oligomerization status. The SARS-CoV-2 N protein is composed of RNA-binding (red) and dimerization domains (green), interspersed with flexible unstructured regions at the N and C-termini and a linker region (yellow) in the middle of the protein. Mutations at three separate sites in the central linker region introduce novel cysteines that differentially increase dimerization levels via a disulfide bond. (RNA-binding: PDB code 6YI3^62^, Dimerization Domain PDB code: 6WZO^23^). We propose that these interactions occur between the linker regions of pairs of N-N dimers and mediate different levels of higher order N-N oligomerization. This figure was made using BioRender.

N is a highly multifunctional protein, and it is likely that some of the regulation of its multiple activities depends on whether it is in the monomeric or oligomerized form. It is thought that phosphorylation of N, specifically within the SR region of the linker, acts as a biological ‘switch’ and mediates the switch between genome packaging/assembly and N’s intracellular roles (including binding NSP3)^8,29–31^. The G215C Delta mutation lies only slightly upstream of key mutations at 203/204 seen in multiple VOCs that modulate viral pathogenicity^8^. It is possible that oligomerization status and phosphorylation states play interacting roles, and that steric considerations affect the ability of cellular kinases to access these residues in the oligomerized state. Nucleocapsid in mature virions is known to be hypo-phosphorylated in the SR region that encompasses both the 203/204 and 215 mutations, in contrast to the intracellular pool which is hyper-phosphorylated^32,33^. Our data indicates that the stably dimerized form of N is packaged at higher levels into mature virions, this effect may be partially due to a decreased ability of cellular kinases to access the key phosphorylation sites.

One of the key roles of the nucleocapsid is to encapsidate the viral RNA during the process of packaging the full-length viral genome into newly forming virions, helping to chaperone and protect the viral genome in its transition from the producer to new target cell. It is known that the nucleocapsid bound to the viral genome is generally in higher order structures (∼14-20 nm, likely composed of 12 copies of N), around which the viral RNA is wrapped (similar to beads on a string)^34–36^. Our data support this framework, as the stably dimerized form of N is seen preferentially in free virus (Fig 6C) compared to transfected or infected cells (Fig 6A,B). Increasing the stability of nucleocapsid oligomers could increase the rate or efficiency of viral encapsidation, which fits with our observation that the G215C mutation increases the amount of nucleocapsid packaged into virions, compared to infectious viral units or other viral structural proteins (Fig 6E,F).

While SARS-CoV-2 virions lack the rigid organization of a virus with a defined icosahedral capsid, a 50-300% increase of nucleocapsid would seem difficult to accommodate within the standard shape a WT virion. Indeed, our data show that, for at least a portion of particles, shifting the oligomerization status by introducing the G215C mutation results in particles that are elongated compared to WT virus (Fig 7). This finding is particularly interesting in light of recent observations that the virions of a clinical Delta isolate retain the spherical shape and radius size found in the ancestral SARS-CoV-2 lineage^37^, suggesting that additional mutations in the Delta lineage have the effect of counteracting the phenotype we observe with the G215C virus.

Relatedly, one of the critical unanswered questions of this study is why no Omicron sub-lineages have possessed such a cysteine in the linker region of the nucleocapsid. It is clear from human transmission data, as well as the clear beneficial effect *in vitro* and *in vivo* (Fig 4E, Fig 5B/C) of the G215C mutation that a cysteine in this region of the nucleocapsid is beneficial. Differing stability of dimer formation at different positions however (greatest at 215>243>185; Fig 2B/3B) suggests that the 215 location is preferred over bond formation closer to either the RNA-binding or dimerization-domains. Furthermore, while the mutations at 214 and 215 were evolutionarily maintained in the Lambda and Delta lineages, the mutations at 243 and 185 we observed in our Iota and Beta stocks were not maintained in these lineages. It is possible that there has not yet been sufficient time for a cysteine-causing mutation to arise at the 214/215 residue of nucleocapsid, and this mutation could occur in a future variant of concern (in the context of omicron or an as-of-yet unknown future variant). Our data suggest this mutation would be likely to increase viral titers significantly, and mutations in this region should be monitored closely for their public health implications. It is also possible that there are epistatic interactions that mediate the beneficial effect of the G215C mutation that are not present in the Omicron background. While the G215C mutation was clearly beneficial for Delta (Fig 1), as well as ancestral SARS-CoV-2 in the absence of any other complementary mutations (Fig 4E, 5B/C), future studies in the Omicron background would be needed to understand the effect this mutation would have on currently circulating virus lineages.

Curiously, the mutations we observe in this study (particularly 215) lie on the binding interface between the N linker and the NSP3 Ubl1 domain^38^, and we hypothesize that stably dimerized N would have a reduced ability to interact with NSP3. There is clear evidence that the interaction between N and NSP3 at this binding interface is important for coronavirus biology, and mismatches between the N linker region and the NSP3 Ubl1 domain severely attenuate viral titers or block viral rescue^39,40^. The function of the N-NSP3 interaction has not been fully resolved, though it has been suggested to play critical roles in either tethering the incoming viral genome to the replication-transcription complex, or delivering the viral genome to the double membrane vesicles (DMVs), to ensure successful early replication^39–41^. N is known to play a key role early in infection, as the gRNA of coronaviruses (unlike nearly all other positive strand RNA viruses) is only minimally infectious in the absence of N protein, suggesting that this N-NSP3 interaction is likely key for efficient replication or transcription of viral RNA^42–46^.

In addition to its role in the replicase, NSP3 forms a key portion of the pore connecting the cytoplasm to the interior of the DMV where viral transcription occurs^47,48^. NSP3 comprises the outermost (cytoplasmic facing) layer of the DMV, with the Ubl1 domain at the very tip of the ‘prongs’ that extend outwards from the pore into the cytoplasm^48^. Recent structural work proposed that the interaction between the linker region of N and the Ubl1 domain of NSP3 helps to mediate a condensation of the nucleocapsid protein prior to encapsidation^38^. In addition, this interaction is proposed to tether nucleocapsid molecules in the immediate vicinity of the DMV exit, ensuring that viral genomes are immediately coated with N as they are extruded out from the DMV pore^38^. We believe that the mutations and viruses characterized in this study may prove useful reagents for future studies studying the functional implications of the N-NSP3 interaction.

Our data raise one additional unanswered question, in that we observe a substantial advantage in viral growth of the N:G215C virus in differentiated human bronchial cells and hamsters but not in Vero E6 cells (Fig 4C vs 4E/6B,C). Vero cells often fail to recapitulate the results seen in human culture systems; they lack the ability to produce and respond to IFN as well as encoding a different suite of cellular proteins, including kinases^3,5,8^. It is possible that the increased oligomerization of N with the G215C mutation drives a greater level of encapsidation that is better able to shield viral RNA from detection by RIG-I and the cellular innate immune response present in primary human lung cells and the hamster model. Alternatively, it is possible that there is a specific host factor present in differentiated lung cells, either a restriction factor that the N dimer is specifically able to evade, or a pro-viral factor that only the dimerized N can access. Future studies using the mutations and viruses characterized in this study will be focused on understanding the mechanism of the growth advantage of N:G215C *in vitro* and *in vivo* and the larger role that levels of N oligomerization play in coronavirus infection.

## MATERALS & METHODS

### Ethics statement

Research conducted in this study was reviewed and approved by the Institutional Biosafety Committee of the University of Vermont (REG202100001) and the Institutional Biosafety Committee of the University of Texas Medical Branch (UTMB). All studies in animals were conducted under a protocol approved by the UTMB Institutional Animal Care and Use Committee and complied with USDA guidelines in a laboratory accredited by the Association for Assessment and Accreditation of Laboratory Animal Care. UTMB is a registered Research Facility under the Animal Welfare Act. It has a current assurance (A3314-01) with the Office of Laboratory Animal Welfare (OLAW), in compliance with NIH Policy. Procedures involving infectious SARS-CoV-2 were performed in the Galveston National Laboratory ABSL3 facility.

The use of deidentified positive clinical specimens was approved by the University of Washington Institutional Review Board (STUDY00000408) and the University of Vermont Institutional Review Board (STUDY00000881) with a consent waiver.

### Cell culture

#### Stable cell lines

Human embryonic kidney cells (HEK-293T/17) (CRL-11268, American Type culture Collection, Manassas, V) were kindly provided by J. Salogiannis (UVM); Vero-E6 cells expressing TMPRSS2 were obtained from the Japanese Cancer Research Resources Bank (JCRB1819). All cell lines were maintained in Dulbecco’s Modified Eagle Medium (DMEM) (10-017-CM, Corning) and supplemented with 10% fetal bovine serum (FBS) (16140-071, Gibco). Cells were grown at 37°C and 5% CO2.

#### Primary cells

24-well transwell inserts (3470-Clear, Costar) were coated with 125uL of a 50ug/mL rat tail collagen (A10483-01, Gibco) in 0.02N acetic acid for 1hr at room temperature. Collagen solution was aspirated and washed once with DMEM (10-017-CM, Corning) before membranes were equilibrated with DMEM in both the apical and basolateral chambers for 1hr at 37°C. DMEM was replaced with “basal media” containing PenumaCult Ex-Plus medium (05040, Stemcell Technologies), 10mL of PneumaCult 50x supplement (05042, StemCell Technologies), 0.001% hydrocortisone (07926, StemCell Technologies), 1% penicillin/streptomycin (30-001-Cl, Corning), 30ug/mL gentamicin (15750060, Gibco) and 15ug/mL amphotericin (30-003-CF, Corning) for 20 mins at 37°C. Expanded human bronchial epithelia cells (CC-2541, Lonza) were plated onto transwell inserts at a density of 40,000 cells/cm2 in warmed basal media. Apical media was changed the next day to remove residual DMSO after which apical and basolateral media was changed every other day until cells were 98-100% confluent. Once at confluency, basal media was aspirated from the apical side and the basolateral chamber media was replaced with PneumaCult-ALI S media (05002, StemCell Technologies), 0.002% heparin (07980, StemCell Technologies), 0.001% hydrocortisone, 1% penicillin/streptomycin, 30ug/mL gentamicin, 10% v/v PneumaCult-ALI supplement (05003, StemCell Technologies), 1% v/v PneumaCult-ALI maintenance Supplement (05006 StemCell Technologies), and 15ug/mL amphotericin. Basolateral chamber media was changed every other day until differentiation occurred, approximately 21 days post airlift.

### Viruses

#### Viral isolation

SARS-CoV-2 WA1/2020 was obtained from the World Reference Center for Emerging Viruses and Arboviruses at the University of Texas Medical Branch. Viral isolates for Alpha (hCoV-19_USA_WA-UWJ17_2021, B.1.1.7.3), Beta (hCoV-19_USA_UWJ01_2021, B.1.351), Gamma (hCoV-19_USA_WA-UWJ05_2021, P.1.17/B.1.1.28.1.17), Delta (hCoV-19_USA_WA-CDC-UW21061761110_2021, AY.44; hCoV-19_USA_WA-UWD03_2021, AY.120.1; hCoV-19_USA_WA-UWD05_2021, AY.3), Epsilon (hCoV-19_USA_WA-UWJ13_2021, B.1.429), Iota (hCoV-19_USA_VT-UVM5927_2021, B.1.526), Mu (hCoV-19_USA_WA-UWM04_2021, B.1.621.1.2/BB.2) and Omicron (hCoV-19/USA/WA-CDC-UW21120651352/2021, BA.1) SARS-CoV-2 variants were obtained from primary clinical specimens. Deidentified nasal swabs were acquired from persons who tested positive for COVID-19 and the variant lineage was determined via Next Generation Sequencing (NGS) or digital droplet PCR either before or after isolation^8^. All isolated specimens were less than eight days old before −80°C storage, stored in saline solution, and had a diagnostic polymerase-chain reaction (PCR) cycle threshold of less than 32. Virus from clinical specimens was isolated in VeroE6-TMPRSS2 cells. Cells were monitored daily for cytopathic effect (CPE) and harvested when ∼50% of the cells exhibited CPE or death. Clarified samples were stored at −80°C and used to generate working viral stocks. Viral stocks were titered by focus forming assay. Next generation sequencing of viral stocks was conducted by the Microbial Genome Sequencing Center or SeqCoast Genomics.

#### Construction of recombinant SARS-CoV-2

The sequence of recombinant wild-type (WT) SARS-CoV-2 is based on the USA-WA1/2020 strain provided by the World Reference Center for Emerging Viruses and Arboviruses (WRCEVA) and originally isolated by the USA Centers for Disease Control and Prevention^49^. Recombinant WT SARS-CoV-2 and mutant viruses were created using a cDNA clone and standard cloning techniques as described previously^24,50^. Construction of WT SARS-CoV-2 and mutant viruses were approved by the University of Texas Medical Branch Biosafety Committee.

### In vitro SARS-CoV-2 infection

#### Vero-TMPRSS2 Cells

Vero-TMPRSS2 cells were seeded in a 24 well plate at 50,000 cells/well one day before infection. The following day, cells were inoculated in 100 ul of virus at the indicated MOI for one hour at 37°C and 5% CO2, after which the inoculum was removed and replaced with fresh growth media. Supernatants were collected at the indicated timepoints, clarified to remove cellular debris and stored at −80°C until time of titering via focus assay.

#### Human Bronchial Epithelial Cells (HBECs)

Differentiated HBECs cells were washed three times with ∼200ul of 37C HEPES buffered saline before being infected with 100ul of virus at an MOI of 0.5 resuspended in ExPneumaCult ALI-S media + supplements (see primary cell maintenance section). After a 1hr incubation at 37°C and 5% CO2, supernatant was removed. The apical side of the cells were washed with 150 ul ExPneumaCult ALI-S media + supplements at the indicated times post infection to collect virus. Viral washes were stored at −80°C until time of titering via focus assay. Basal media was changed every 48hrs during the infection time course.

### Focus Forming Assay

Viral titrations were performed largely as previously described^51^. VeroE6-TMPRSS2 cells were seeded at a density of 60,000 cells/well into white 96-well plates (Falcon; #353296). 24hrs later, viral samples of interest were serially diluted 10-fold in DMEM + 10% FBS. Plates were aspirated and infected with diluted viral samples for 1hr at 37°C and 5% CO2 before being overlaid with a 1.2% methylcellulose (Acros; #332620010) solution suspended in DMEM. After 24hrs, the methylcellulose solution was aspirated, and the plates were fixed a 4% formaldehyde solution (Honeywell; #F1635-4L) for 20mins before being washed once with deionized (DI) water. Cells were permeabilized in 50ul of 0.05% Triton X100 (Fisher; #BP151-100) for 5mins, washed with PBS and blocked (5% non-fat milk solution in PBS) for 1hr at room temperature.

Virally infected cells were detected with an anti-SARS-CoV-2 N antibody (Sinobiological; #40143-R001, 1:20,000) resuspended in 5% milk for 1hr at 37°C. Cells were washed twice with PBS and stained with a HRP-conjugated secondary antibody (Seracare; #5220-0337, 1:4,000) resuspended in 5% milk for 1hr at 37°C. Cells were washed twice with PBS, and foci developed using a TruBlue HRP substrate (SeraCare; #5510-0030). Foci were imaged on a BioTek ImmunoSpot S6 MACRO Analyzer and manually counted.

### Viral Concentration

Viral stocks were concentrated by vortexing 1 ml of high titer viral stocks with 10% polyethylene glycol (PEG) (Sigma-Aldrich; #P6667) for ∼5-10 minutes before centrifugation at 10,000g for 30mins at 4°C. Pellets were then resuspended in a NP-40, 1% Trition X-100 lysis buffer (see SDS-PAGE and western blotting section below).

### Analysis of publicly available genomic data

The identity of the amino acid residue in the nucleocapsid protein (including at 215) in 7,342,041 SARS-CoV-2 sequences deposited in GenBank, the China National Center for Bioinformation and from COG-UK were analyzed using the phylogenetic tree (Cov2Tree) and visualized by Taxonium on January 10^th^, 2024^52,53^. Sequences for non-SARS-CoV-2 coronavirus nucleocapsid sequences were obtained through the NIH Protein Databank and aligned using Clustal Omega (EMBL-EBI, Clustal 0 (1.2.4))^54^.

### Hamster infection studies

For *in vivo* studies, three- to four-week-old male hamsters were purchased from Envigio, and all studies conducted within the Galveston National Laboratory ABSL3 facility. Studies were conducted in accordance with a protocol approved by the University of Texas Medical Branch Institutional Animal Care and Use Committee and comply with the United States Department of Agriculture guidelines. All laboratories were accredited by the Association for Assessment and Accreditation of Laboratory Animal Care. For infection studies, animals housed in groups of five were intranasally infected PBS alone (mock) or with 10^4^ FFU of WT or N:G215C mutant viruses. Animals were then monitored daily for weight loss and development of clinical disease until completion of the experiment. All procedures were carried out under anesthesia with isoflurane (Henry Schein Animal Health), with the exception of weighing.

### Histology

Hematoxylin- and eosin-stained microscopic slides of lung from each hamster were separated into three groups according to the number of days after intranasal inoculation of SARS Cov 2. Microscopic slides of each group were scrambled thoroughly and examined blindly to assess the severity of pathologic lesions. Slides were placed in the order of serial pairwise comparison of the extent of lesions. Upon completion of the ordering of severity, numbers were assigned to the slides from 1, least pathologic change, to the highest number for the slide with the most severe pathology. At this point the code was revealed, and the sum of the rank order of severity numbers for each group: mock, wild type, and mutant was calculated and divided according to the number of slides in the group to determine a severity score for the group. The number of slides in the groups varied slightly as tissues which contained abundant polymorphonuclear leukocytes, very likely indicating superimposed bacterial bronchopneumonia were removed and not scored. Separate sums of rank order scores and average severity score were calculated for each day.

### Antigen Staining

Antigen staining was performed as previously described^55^. Briefly, cut sections were deparaffinized by heading at 56°C for 10 minutes, followed by 3 washes with xylene and 4 washes with graded ethanol. Slides were then rehydrated with distilled water and antigen retrieval performed by steaming slides for 40 minutes in antigen retrieval solution (10 mM sodium citrate, 0.05% Tween-20, ph6). After cooling, slides were rinsed with water, and endogenous peroxidases quenched in TBS + 0.3% hydrogen peroxide. Sections were then washed with TBST, blocked with 10% goat serum diluted in 1% BSA/TBST, followed by probing with primary anti-N antibody (Sino #40143-R001) at 1:1000. Sections were then washed in 3 times with TBST and incubated in secondary HRP-conjugated anti-rabbit antibody (Cell Signaling Technology #7074) at 1:200. To visualize antigen, sections were treated with ImmPACT NovaRED (Vector Laboratories #SK-4805) for 3 minutes. Sections were then counter stained with hematoxylin for 5 min. Finally, sections were the dehydrated by incubation in graded ethanol followed by xylene and mounted with cover slips. Viral antigen staining occurred through blinded scoring on a scale of 0 to 4, with the scores of two lung sections being averaged to determine the final score.

### Plasmid Generation

The sequences for the N protein of our Beta, Delta, and Iota SARS-CoV-2 variants, as well as the WA1/2020 (‘WT’) sequences were synthesized and subcloned into a pcDNA3.1 vector (GenScript). In addition, point mutants were generated in which the cysteine in the Beta, Iota, and Delta N sequences was mutated back to the corresponding residue in WA1 (Beta: C185R, Delta: C215G, Iota:C243G) or where cysteine were introduced at the 185, 215 or 243 positions into the WA1 background (R185C, G215C, G243C). Plasmid sequences were confirmed via whole-plasmid sequencing (Plasmidsaurus).

### Transfection

HEK-293T cells were seeded at a density of 350,000 cells/well in a 12-well plate (Corning #3512). The following day, 500 ng of the appropriate plasmid and 2μl of Lipofectamine (Invitrogen #52887) diluted to a volume of 40ul in DMEM were incubated for 24hrs at 37°C and 5% CO2. Cells were harvested by scraping into PBS, pelleted and lysed for SDS-PAGE and western blotting (see below).

### SDS-PAGE and western blotting

Cells were lysed in a NP-40 lysis buffer (Thermo Scientific #J60766.AK) containing a 1% Triton X-100 solution (Fisher, #BP151-100) and protease inhibitors (Thermo Scientific Fisher #A32955) for 20mins on ice. Cell lysates were clarified by spinning at 14,000rpm for 10mins at 4**°**C to remove insoluble debris and diluted in a 1:1 ratio (v/v) of 4X Laemmli sample buffer (250 mM Tris-HCl pH 6.8,40% glycerol, 8% SDS, 0.04% Bromophenol Blue). Reduced lysates were treated with either 2.75mM 2-mercaptoethanol (Gibco #21985-023) or 10mM dithiothreitol (Fisher Bioreagents # BP172-5). For NEM processing, samples were lysed in 1.25 M n-ethylmaleimide (NEM; Thermo Fisher #23030 and incubated at 4°C overnight. All samples were boiled for 5 mins before loading. Samples were separated in NuPAGE 4-12% Bis-Tris gels (Invitrogen # NPO335BOX) in MES buffer (Invitrogen # NP0002) with a molecular mass ladder (Thermo Fisher #LC5925) at 180v for 50mins before being transferred into a nitrocellulose membrane (Invitrogen # IB23001) using an iBlot 2 machine (Invitrogen) at 20V for 7mins.

Membranes were blocked in a 5% milk/PBS solution for 30mins and then incubated overnight at 4**°**C in a solution containing the primary antibody in a 5% milk/PBST (0.2% Tween 20 (Thermo Fisher Scientific, # J20605-AP). Membranes were washed in PBST, incubated while rocking with secondary antibodies diluted in 5% milk/PBST for 45 min, washed again with PBST and imaged with a LI-COR Odyssey CLx. Protein expression was analyzed by measuring band densitometry in Fiji (22743772).

### Antibodies

| **Primary Antibodies** |  |  |
| --- | --- | --- |
| **Species, Target** | **Catalog #, Company** | **Dilution Used** |
| SARS-CoV-2 nucleocapsid protein (mouse) | Invitrogen, MA5-35943 | 1:1,000 |
| SARS-CoV-2 nucleocapsid protein (rabbit) | SinoBiological, 40143-R001 | 1:10,000 (western)<br>1:20,000 (focus assay) |
| SARS-CoV-2 membrane (M) protein (mouse) | Cell Signaling, E5A8A | 1:1000 |
| β-actin (mouse) | Novus Biologicals, NB600-501 | 1:10,000 |
| **Secondary Antibodies** |  |  |
| IRDye 680LT goat anti-rabbit IgG | LICOR, #92668021 | 1:10,000 |
| IRDye 680RD goat anti-mouse IgG | LICOR, #92668070 | 1:10,000 |
| IRDye 800CW goat anti-mouse IgG | LICOR, #92632210 | 1:10,000 |
| HRP anti-rabbit | Seracare; #5220-0337 | 1:4,000 |

#### Immunoprecipitation assays

VeroE6-TMPRSS2 cells were infected at an MOI of 0.01 with WT (WA1) or Delta SARS-CoV-2. 24hpi cells were scraped into PBS, pelleted and lysed in NP-40/Triton lysis buffer (see SDS-PAGE and western blotting. SARS-CoV-2 N was affinity purified using 10 ul of an anti-SARS-CoV-2 N antibody (40143-R001, Sinobiological) and Protein G-conjugated Dynabeads (Invitrogen, #10003D) according to the manufacturer instructions.

#### Mass Spectrometry

After immunoprecipitation, samples were run on an 8% BOLT gel (Invitrogen, # NW00085BOX), bands were visualized via Coomassie stain (40% methanol, 20% acetic acid, 0.01% Brilliant Blue) and de-stained (30% methanol, 10% acetic acid). Bands of interest (monomer, dimer, or full lane) were cut out of the gel, and further cut into ∼1 mm cubes before being placed into tubes for processing using HPLC-grade Fisher brand chemicals. Gel slices were incubated in water for 5 min, and then de-stained for 30 min at 37 **°**C (50 mM ammonium bicarbonate, 50% acetonitrile). Slices were rinsed in fresh acetonitrile for two 5 min incubations before incubating for 30 min at 55 **°**C in 25 mM DTT (Thermo Fisher, #R0861)/50 mM ammonium biocarbonate.

Slices were cooled, incubated in 100% acetonitrile for 5 min and incubated for 45 min at room temperature in 10 mM iodoacetamide (Sigma-Aldrich, #I6125)/50 mM ammonium bicarbonate, in the dark. Slices were incubated in (50 mM ammonium bicarbonate, 50% acetonitrile) for 5 min, rinsed in water for 10 min and centrifuged at 13,000 rpm for 30 s. Next, samples were dehydrated in acetonitrile before all liquid removed and gel slices allowed to dry completely.

Once dry, samples were placed on ice for 5 min before incubating for 30 min on ice with 6 ng/µl trypsin (Promega #V511A) in 50 mM ammonium bicarbonate and then digesting overnight at 37 **°**C. Supernatants were removed and saved, while gel slices were covered with a 50% acetonitrile, 2.5% formic acid solution and spun for ten min after which supernatants were combined with those from the previous step. Finally, 30 µl acetonitrile was added to gel slices for a final 10 min incubation, after which these supernatants were combined with those from the previous two steps and dried in a vacuum centrifuge at 37 **°**C.

Following peptide preparation, samples were resuspended in 2.5% acetonitrile and 0.1% formic acid. Then loaded onto an EASY n-LC 1200 for HPLC (300 nL/min) followed by tandem MS/MS on an Exploris mass spectrometer (ThermoFisher). The Exploris was fitted with a Nanospray Flex ion source and chromatograph column packed in-house (35 cm x 100 µm, 1.8 µm 120 Å UChrom, C18 packing material). Peptides were eluted into the mass spectrometer on a 5-95% gradient of 80% acetonitrile and 0.1% formic acid over 170 minutes, followed by a 10-min clear at 5% gradient. The following were the mass spectrometer parameters. Precursor scan range = 350-1400 *m/z*, resolution = 60,000, normalized AGC target = 250% of maximum, maximum IT = 25 ms. Data-dependent MS2 orbitrap resolution = 15,000, normalized AGC target = 50% of maximum, isolation window = ± 1.4 *m/z*, collision energies = 30%, dynamic exclusion = 45 s with repeat count of 1. Followed by ten collision induced dissociation tandem mass spectra of the top ten most abundance ions in the precursor scan.

Resulting raw mass spectra for each sample was searched against both a forward and reverse African Green Monkey protein database and a SARS CoV-2 protein database using SEQUEST^56^. Search parameters included a requirement for tryptic peptides and differential modifications (phosphorylation on serine, threonine, and tyrosine (+79.9663 Da), oxidation of methionine (+15.99.49 Da), acrylamindation of cysteine (+71.0371 Da), and carboxyamidomethylation of cysteine (+57.0214 Da)). The resulting peptide lists were filtered by mass accuracy (ppm ≤± 5), cross correlation scores (*z*=1 XCorr ≥ 1.8, *z*=2 XCorr ≥ 2, *z*=3 XCorr ≥ 2.2*, z*=4 XCorr ≥ 2.4, *z*=5 XCorr ≥ 2.6), and a unique delta-correlation (uniqΔcorr ≥ 0.15). All filtering resulted in a false discovery rate of less than 0.01%. The filtered peptide lists were then compared by sample type (present or absent or enriched (2.5-fold increase)) by spectral count using R software.

#### Sample preparation for electron microscopy

Vero-E6 cells were cultured on plates and infected with mNG WT or G215C SARS-CoV-2 under BSL-3 conditions. Cells were pre-fixed with 3% glutaraldehyde, 1% paraformaldehyde, 5% sucrose in 0.1 M sodium cacodylate, then removed from the plates and transferred to Eppendorf tubes and gently pelleted. The fixative supernatant was removed and pellets rinsed with fresh cacodylate buffer containing 10% Ficoll, placed into brass planchettes (Ted Pella, Inc.), and rapidly frozen with an HPM-010 high-pressure freezing machine (Bal-Tec, Leichtenstein). The frozen samples were transferred under liquid nitrogen to cryotubes (Nunc) containing a frozen solution of 2.5% osmium tetroxide, 0.05% uranyl acetate in acetone. Tubes were loaded into an AFS-2 freeze-substitution machine (Leica Microsystems, Vienna) and processed at −90°C for 72 h, warmed over 12 h to −20°C, held at that temperature for 6 h, then warmed to 4°C for 1 h. Samples were rinsed 3x with cold acetone, after which they were infiltrated with Epon-Araldite resin (Electron Microscopy Sciences) over 48 h. Cell pellets were flat-embedded between two Teflon-coated glass microscope slides and the resin was polymerized at 60°C for 48 h.

#### Electron microscopy and Dual-Axis tomography

Embedded cells were observed by light microscopy and appropriate blocks were extracted with a microsurgical scalpel and glued to the tips of plastic sectioning stubs. Semi-thin (170 nm) serial sections were cut with a UC6 ultramicrotome (Leica Microsystems) using a diamond knife (Diatome Ltd., Switzerland). Sections were placed on formvar-coated copper-rhodium slot grids (Electron Microscopy Sciences) and stained with 3% uranyl acetate and lead citrate. Gold beads (10 nm) were placed on both surfaces of the grid to serve as fiducial markers for subsequent image alignment. Grids were placed in a dual-axis tomography holder (Model 2040, E.A. Fischione Instruments) and imaged with a Tecnai T12-G2 transmission electron microscope operating at 120 KeV (ThermoFisher Scientific) equipped with a 2k x 2k CCD camera (XP1000; Gatan, Inc.). Tomographic tilt-series and large-area montaged overviews were acquired automatically using the SerialEM software package^57^. For tomography, samples were tilted ±62° and images collected at 1° intervals. The grid was then rotated 90° and a similar series taken about the orthogonal axis. Tomographic data was calculated, analyzed, and modeled using the IMOD software package^58–60^ on iMac Pro and Mac Studio M1 computers (Apple, Inc.).

#### Statistics

Unpaired T-tests on raw or log-transformed (viral titer) data were performed using GraphPad Prism 10 or R^61^. Statistical significance is indicated with. (p=.05-.1), * (p<0.05), ** (p<0.01), *** (p<.001) or **** (p<.0001).

## Data Availability

All raw sequencing data are available in the NCBI Bioproject ID PRJNA1083584. Individual peptides identified in proteomics experiments are included in supplementary table 1, raw spectrometry files are available upon request.

## Resources

The authors thank the Caltech Beckman Institute CryoEM Facility for use and maintenance of the Tecnai T12 TEM.

## Acknowledgements

We would like to thank the University of Vermont Proteomics Core Facility, Drs Matthew Poynter, Dimitry Krementsov and Joyce Oetjen for technical assistance.

## Author contributions

Conceptualization (HCK, HWD, EAB), Formal Analysis (HCK, HWD, BAJ, CMD, DJS, BAB, VDM, EAB), Investigation (HCK, HWD, BAJ, MMS, SJ, KL, CMD, DHW, BAB, MSL), Reagents

(MGM, AP, JWC, PR, ALG, KRJ, VDM), Writing-Original Draft (HCK, HWD, EAB), Writing-Review & Editing (All authors), Visualization (HCK, HWD, BAJ, MSL, EAB), Supervision (BAB, PJB, VDM, EAB), Funding Acquisition (VDM, EAB).

---

**Supplementary Figure 1:**
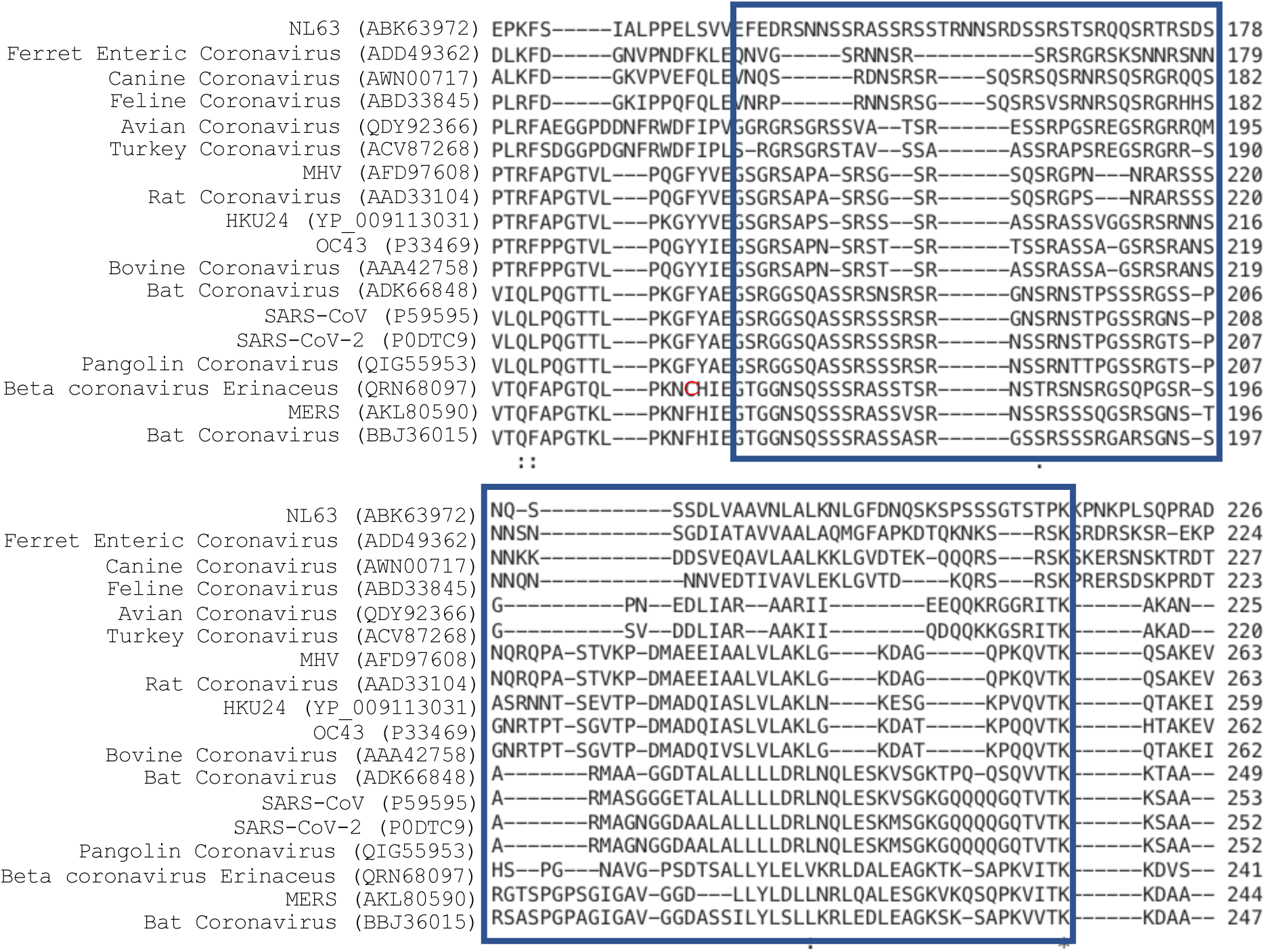
Absence of cysteines in the linker region of Coronaviridae nucleocapsid proteins. Sequences for the indicated coronavirus nucleocapsid protein sequences were obtained through the NIH Protein Databank and aligned using Clustal Omega from EMBL-EBI (Clustal 0 (1.2.4)). The navy box highlights the linker region (residues 175-247 in SARS-CoV-2). Shown in red is the only cysteine in the displayed sequences, which occurs immediately before the linker region in Erinaceus betacoronavirus.

**Supplementary Figure 2:**
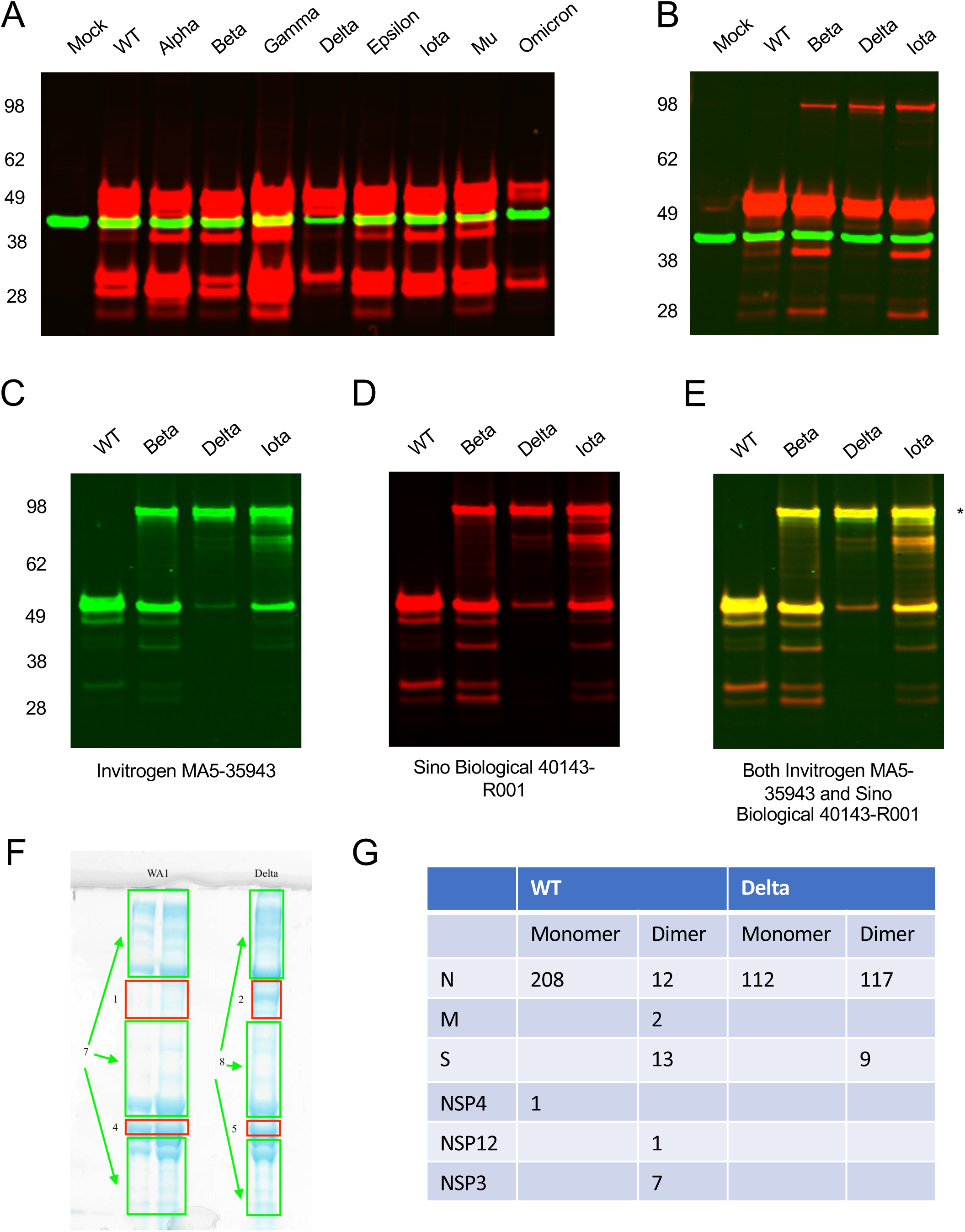
SARS-CoV-2 N dimer formation. VeroE6-TMPRSS2 cells were infected, or mock infected, with the indicated SARS-CoV-2 variants (or WA1, termed Wildtype) at an MOI of 0.01 for 24 hours. Cell lysates were harvested under A) reducing conditions (10 mM DTT) or B) non-reducing conditions, but in the presence of NEM (which prevents the formation of post-lysis disulfide bonds) before visualization by SDS-PAGE and western blot using antibodies recognizing SARS-CoV-2 N (red) and β-actin (green). Alternatively, unreduced lysates from cells infected as above were visualized by SDS-PAGE and western blot using two different antibodies recognizing SARS-CoV-2 N: (C) Invitrogen anti-N mouse antibody (MA5-35943, in green) and (D) Sino Biological anti-N rabbit antibody (40143-R001 in red). (E) Lysates were stained and imaged simultaneously with the two anti-N antibodies and the overlay is shown in yellow. Note minor bands that are seen with the Invitrogen but not the Sino-Biological antibody. The MW of the ∼100 kDa band observed in Beta, Delta and Iota samples under non-reducing conditions is indicated by a *. A representative gel from 3 (A, B) or 2 (C) independent biological replicates is shown. F) Vero-TMPRSS2 cells were infected with WA1 or Delta SARS-CoV-2 at an MOI of 0.01. 24 hpi cells were harvested, lysed and N was affinity purified using immunoprecipitation. N and associated proteins were run on an SDS-PAGE gel, and bands corresponding to the monomer and dimer were cut and processed for mass spectrometry. G) The number of peptides for each SARS-CoV-2 viral protein found in the monomer or dimer portion of the gel, for either WT (WA1) or Delta, is shown.

**Supplementary Figure 3:**
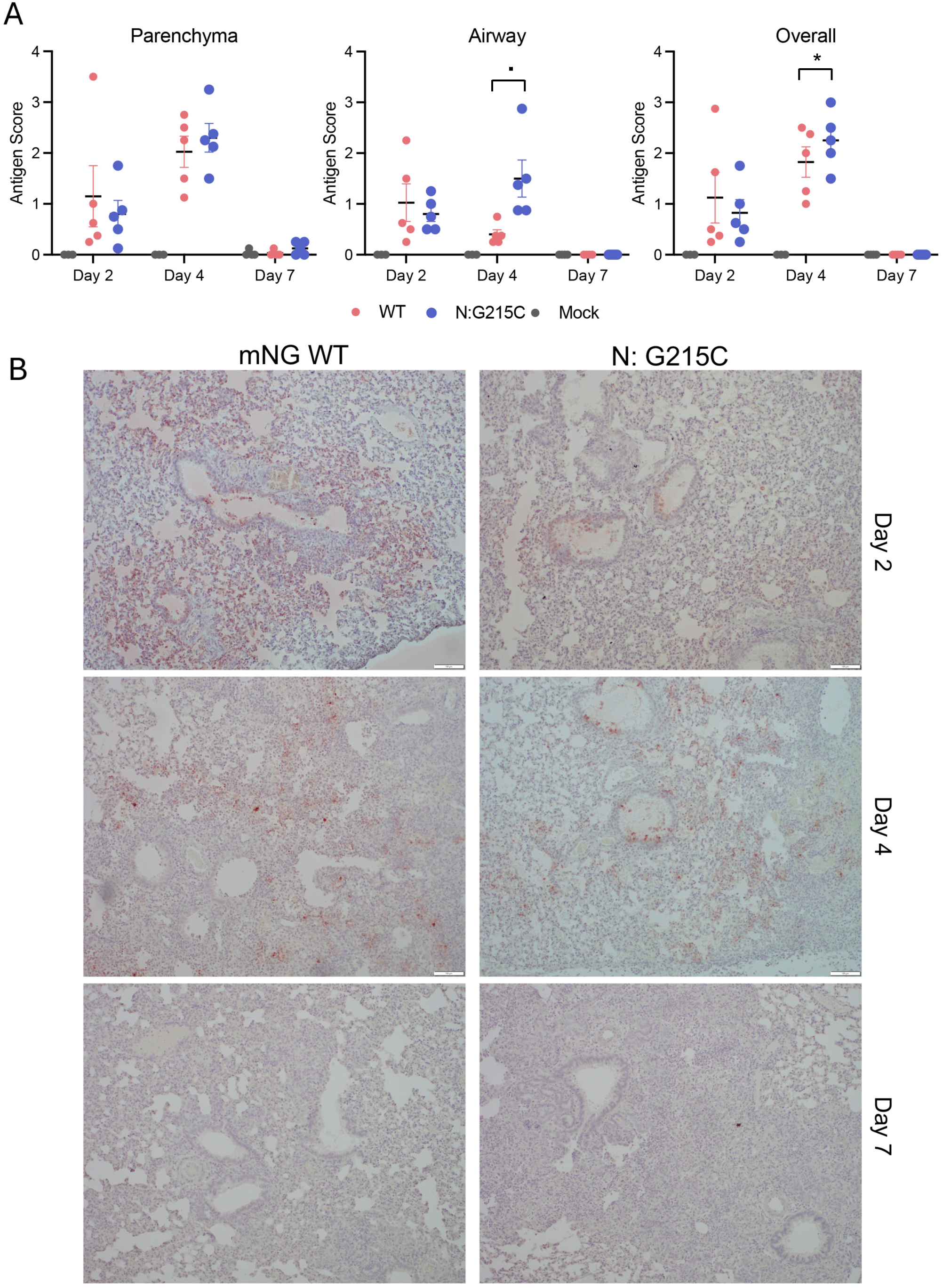
Antigen staining of lung tissue from infected hamsters. Section of lung tissue from infected hamsters were stained for viral antigen (nucleocapsid). Sections were then blinded and scored on a 4 point scale for parenchyma, airway, and overall staining (A). Each data point represents the average score form two lung section from each hamster in the group (n>4). Horizontal lines represent the group mean and error bars are +/- SD. Significance determined by student’s T-test. (B) Representative images are shown for WT and N:215C infected animals for 2, 4, and 7 dpi. [. (p=.05-.1), * (p<0.05)].

**Supplementary Figure 4:**
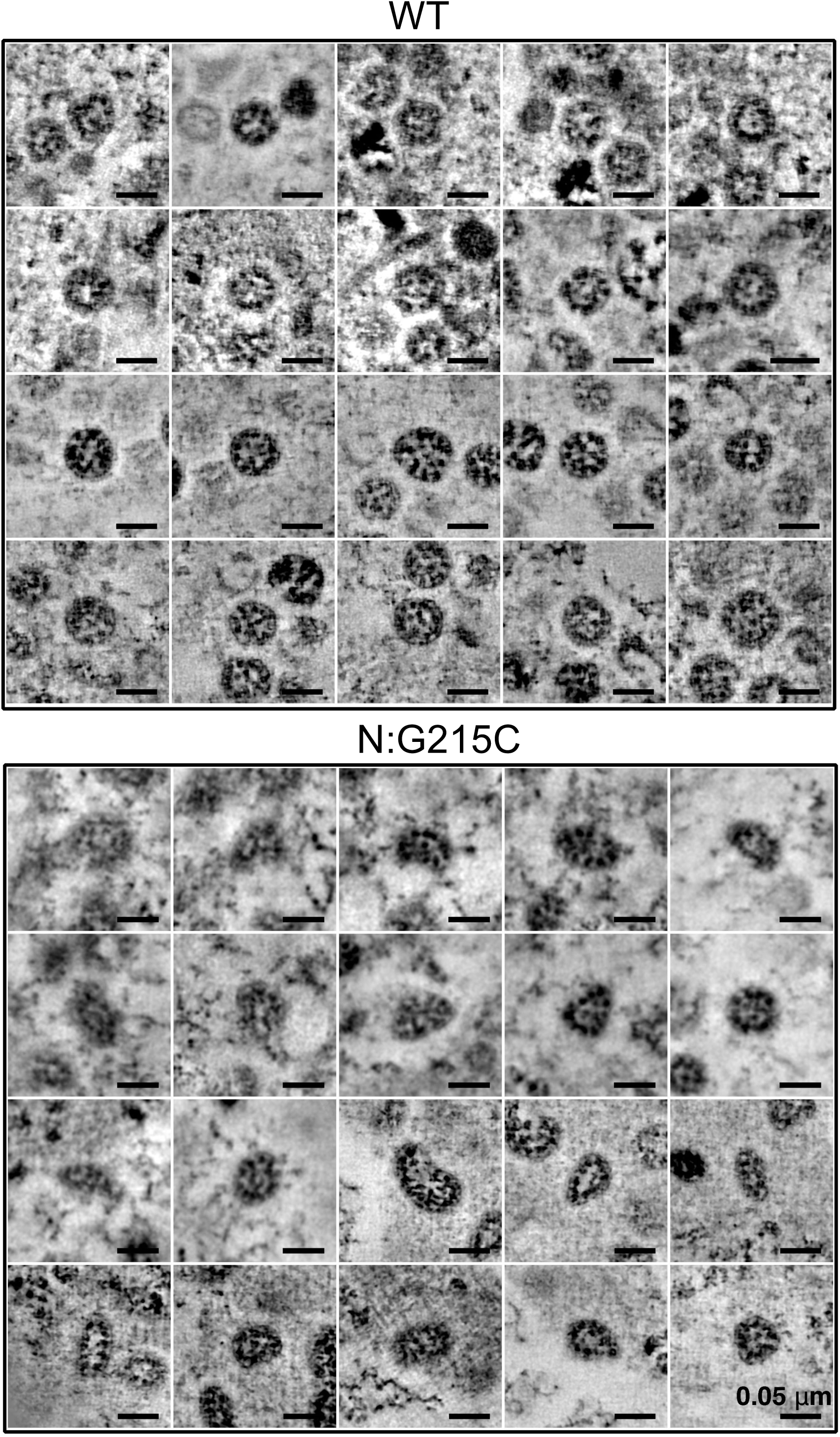
N:G215C virions are enlarged. Vero-TMPRSS2 cells were infected with WT or N:G215C viruses at an MOI of 0.1. The following day cells were prepared for electron microscopy by high-pressure freezing and freeze-substitution, then sectioned and imaged by dual-axis electron tomography. Virus-containing exit compartments were located in both samples, and 20 virions that had completely separated from cellular membranes were randomly selected and imaged for each virus.

## References Cited

1. Cucinotta, D. & Vanelli, M. WHO Declares COVID-19 a Pandemic. Acta Biomed. Atenei Parm. 91, 157–160 (2020).

2. Plante, J. A. et al. Spike mutation D614G alters SARS-CoV-2 fitness. Nature 592, 116–121 (2021).

3. Johnson, B. A. et al. Loss of furin cleavage site attenuates SARS-CoV-2 pathogenesis. Nature 591, 293–299 (2021).

4. Liu, Y. et al. Delta spike P681R mutation enhances SARS-CoV-2 fitness over Alpha variant. Cell Rep. 39, (2022).

5. Vu, M. N., et al. QTQTN motif upstream of the furin-cleavage site plays a key role in SARS-CoV-2 infection and pathogenesis. Proc. Natl. Acad. Sci. 119, e2205690119 (2022).

6. McGrath, M. E., et al. SARS-CoV-2 variant spike and accessory gene mutations alter pathogenesis. Proc. Natl. Acad. Sci. 119, e2204717119 (2022).

7. Carabelli, A. M. et al. SARS-CoV-2 variant biology: immune escape, transmission and fitness. Nat. Rev. Microbiol. 21, 162–177 (2023).

8. Johnson, B. A. et al. Nucleocapsid mutations in SARS-CoV-2 augment replication and pathogenesis. PLOS Pathog. 18, e1010627 (2022).

9. Chang, C., Hou, M.-H., Chang, C.-F., Hsiao, C.-D. & Huang, T. The SARS coronavirus nucleocapsid protein – Forms and functions. Antiviral Res. 103, 39–50 (2014).

10. Zúñiga, S. et al. Coronavirus Nucleocapsid Protein Facilitates Template Switching and Is Required for Efficient Transcription. J. Virol. 84, 2169–2175 (2010).

11. Almazán, F., Galán, C. & Enjuanes, L. The Nucleoprotein Is Required for Efficient Coronavirus Genome Replication. J. Virol. 78, 12683–12688 (2004).

12. Schelle, B., Karl, N., Ludewig, B., Siddell, S. G. & Thiel, V. Selective Replication of Coronavirus Genomes That Express Nucleocapsid Protein. J. Virol. 79, 6620–6630 (2005).

13. Thiel, V., Herold, J., Schelle, B. & Siddell, S. G. Viral Replicase Gene Products Suffice for Coronavirus Discontinuous Transcription. J. Virol. 75, 6676–6681 (2001).

14. Hurst, K. R. et al. A Major Determinant for Membrane Protein Interaction Localizes to the Carboxy-Terminal Domain of the Mouse Coronavirus Nucleocapsid Protein. J. Virol. 79, 13285–13297 (2005).

15. Verma, S., Bednar, V., Blount, A. & Hogue, B. G. Identification of Functionally Important Negatively Charged Residues in the Carboxy End of Mouse Hepatitis Coronavirus A59 Nucleocapsid Protein. J. Virol. 80, 4344–4355 (2006).

16. Kuo, L., Hurst-Hess, K. R., Koetzner, C. A. & Masters, P. S. Analyses of Coronavirus Assembly Interactions with Interspecies Membrane and Nucleocapsid Protein Chimeras. J. Virol. 90, 4357–4368 (2016).

17. Dutta, N. K., Mazumdar, K. & Gordy, J. T. The Nucleocapsid Protein of SARS–CoV-2: a Target for Vaccine Development. J. Virol. 94, 10.1128/jvi.00647-20 (2020).

18. Liu, Y. et al. The N501Y spike substitution enhances SARS-CoV-2 infection and transmission. Nature 602, 294–299 (2022).

19. Saito, A. et al. Enhanced fusogenicity and pathogenicity of SARS-CoV-2 Delta P681R mutation. Nature 602, 300–306 (2022).

20. Zhao, H., et al. Plasticity in structure and assembly of SARS-CoV-2 nucleocapsid protein. PNAS Nexus 1, pgac049 (2022).

21. Chu, H. et al. Coronaviruses exploit a host cysteine-aspartic protease for replication. Nature 609, 785–792 (2022).

22. Zhao, H. et al. A conserved oligomerization domain in the disordered linker of coronavirus nucleocapsid proteins. Sci. Adv. 9, eadg6473 (2023).

23. Ye, Q., West, A. M. V., Silletti, S. & Corbett, K. D. Architecture and self-assembly of the SARS-CoV-2 nucleocapsid protein. Protein Sci. 29, 1890–1901 (2020).

24. Xie, X. et al. An Infectious cDNA Clone of SARS-CoV-2. Cell Host Microbe 27, 841–848.e3 (2020).

25. Jack, A. et al. SARS-CoV-2 nucleocapsid protein forms condensates with viral genomic RNA. PLOS Biol. 19, e3001425 (2021).

26. Aloise, C. et al. SARS-CoV-2 nucleocapsid protein inhibits the PKR-mediated integrated stress response through RNA-binding domain N2b. PLOS Pathog. 19, e1011582 (2023).

27. LeBlanc, K. et al. The Nucleocapsid Proteins of SARS-CoV-2 and Its Close Relative Bat Coronavirus RaTG13 Are Capable of Inhibiting PKR- and RNase L-Mediated Antiviral Pathways. Microbiol. Spectr. 11, e00994–23 (2023).

28. Liu, H. et al. SARS-CoV-2 N Protein Antagonizes Stress Granule Assembly and IFN Production by Interacting with G3BPs to Facilitate Viral Replication. J. Virol. 96, e00412–22 (2022).

29. Cubuk, J. et al. The SARS-CoV-2 nucleocapsid protein is dynamic, disordered, and phase separates with RNA. Nat. Commun. 12, 1936 (2021).

30. Lu, S. et al. The SARS-CoV-2 nucleocapsid phosphoprotein forms mutually exclusive condensates with RNA and the membrane-associated M protein. Nat. Commun. 12, 502 (2021).

31. Carlson, C. R. et al. Phosphoregulation of Phase Separation by the SARS-CoV-2 N Protein Suggests a Biophysical Basis for its Dual Functions. Mol. Cell 80, 1092–1103.e4 (2020).

32. Wu, C.-H. et al. Glycogen Synthase Kinase-3 Regulates the Phosphorylation of Severe Acute Respiratory Syndrome Coronavirus Nucleocapsid Protein and Viral Replication *. J. Biol. Chem. 284, 5229–5239 (2009).

33. Wu, C.-H., Chen, P.-J. & Yeh, S.-H. Nucleocapsid Phosphorylation and RNA Helicase DDX1 Recruitment Enables Coronavirus Transition from Discontinuous to Continuous Transcription. Cell Host Microbe 16, 462–472 (2014).

34. Yao, H. et al. Molecular Architecture of the SARS-CoV-2 Virus. Cell 183, 730–738.e13 (2020).

35. Klein, S. et al. SARS-CoV-2 structure and replication characterized by in situ cryo-electron tomography. Nat. Commun. 11, 5885 (2020).

36. Bracquemond, D. & Muriaux, D. Betacoronavirus Assembly: Clues and Perspectives for Elucidating SARS-CoV-2 Particle Formation and Egress. mBio 12, 10.1128/mbio.02371-21 (2021).

37. Ke, Z. et al. Virion morphology and on-virus spike protein structures of diverse SARS-CoV-2 variants. 2023.12.21.572824 Preprint at 10.1101/2023.12.21.572824 (2023).

38. Bessa, L. M. et al. The intrinsically disordered SARS-CoV-2 nucleoprotein in dynamic complex with its viral partner nsp3a. Sci. Adv. 8, eabm4034 (2022).

39. Hurst, K. R., Koetzner, C. A. & Masters, P. S. Characterization of a Critical Interaction between the Coronavirus Nucleocapsid Protein and Nonstructural Protein 3 of the Viral Replicase-Transcriptase Complex. J. Virol. 87, 9159–9172 (2013).

40. Hurst, K. R., Ye, R., Goebel, S. J., Jayaraman, P. & Masters, P. S. An Interaction between the Nucleocapsid Protein and a Component of the Replicase-Transcriptase Complex Is Crucial for the Infectivity of Coronavirus Genomic RNA. J. Virol. 84, 10276–10288 (2010).

41. Koetzner, C. A., Hurst-Hess, K. R., Kuo, L. & Masters, P. S. Analysis of a crucial interaction between the coronavirus nucleocapsid protein and the major membrane-bound subunit of the viral replicase-transcriptase complex. Virology 567, 1–14 (2022).

42. Yount, B. et al. Reverse genetics with a full-length infectious cDNA of severe acute respiratory syndrome coronavirus. Proc. Natl. Acad. Sci. 100, 12995–13000 (2003).

43. Yount, B., Denison, M. R., Weiss, S. R. & Baric, R. S. Systematic Assembly of a Full-Length Infectious cDNA of Mouse Hepatitis Virus Strain A59. J. Virol. 76, 11065–11078 (2002).

44. Yount, B., Curtis, K. M. & Baric, R. S. Strategy for Systematic Assembly of Large RNA and DNA Genomes: Transmissible Gastroenteritis Virus Model. J. Virol. 74, 10600–10611 (2000).

45. Coley, S. E. et al. Recombinant Mouse Hepatitis Virus Strain A59 from Cloned, Full-Length cDNA Replicates to High Titers In Vitro and Is Fully Pathogenic In Vivo. J. Virol. 79, 3097–3106 (2005).

46. Casais, R., Thiel, V., Siddell, S. G., Cavanagh, D. & Britton, P. Reverse Genetics System for the Avian Coronavirus Infectious Bronchitis Virus. J. Virol. 75, 12359–12369 (2001).

47. Snijder, E. J. et al. A unifying structural and functional model of the coronavirus replication organelle: Tracking down RNA synthesis. PLOS Biol. 18, e3000715 (2020).

48. Wolff, G. et al. A molecular pore spans the double membrane of the coronavirus replication organelle. Science 369, 1395–1398 (2020).

49. Harcourt, J. et al. Severe Acute Respiratory Syndrome Coronavirus 2 from Patient with Coronavirus Disease, United States - Volume 26, Number 6—June 2020 - Emerging Infectious Diseases journal - CDC. doi:10.3201/eid2606.200516.

50. Xie, X. et al. Engineering SARS-CoV-2 using a reverse genetic system. Nat. Protoc. 16, 1761–1784 (2021).

51. Despres, H. W. et al. Measuring infectious SARS-CoV-2 in clinical samples reveals a higher viral titer:RNA ratio for Delta and Epsilon vs. Alpha variants. Proc. Natl. Acad. Sci. 119, e2116518119 (2022).

52. Sanderson, T. Taxonium, a web-based tool for exploring large phylogenetic trees. eLife 11, e82392 (2022).

53. Kramer, A. M., Sanderson, T. & Corbett-Detig, R. Treenome Browser: co-visualization of enormous phylogenies and millions of genomes. Bioinformatics 39, btac772 (2023).

54. Madeira, F. et al. Search and sequence analysis tools services from EMBL-EBI in 2022. Nucleic Acids Res. 50, W276–W279 (2022).

55. Garvanska, D. H. et al. The NSP3 protein of SARS-CoV-2 binds fragile X mental retardation proteins to disrupt UBAP2L interactions. EMBO Rep. 25, 902–926 (2024).

56. An approach to correlate tandem mass spectral data of peptides with amino acid sequences in a protein database. - University of Vermont. https://primo.uvm.edu.

57. Mastronarde, D. N. Automated electron microscope tomography using robust prediction of specimen movements. J. Struct. Biol. 152, 36–51 (2005).

58. Kremer, J. R., Mastronarde, D. N. & McIntosh, J. R. Computer Visualization of Three-Dimensional Image Data Using IMOD. J. Struct. Biol. 116, 71–76 (1996).

59. Mastronarde, D. N. Correction for non-perpendicularity of beam and tilt axis in tomographic reconstructions with the IMOD package. J. Microsc. 230, 212–217 (2008).

60. Mastronarde, D. N. & Held, S. R. Automated tilt series alignment and tomographic reconstruction in IMOD. J. Struct. Biol. 197, 102–113 (2017).

61. R: The R Project for Statistical Computing. https://www.r-project.org/.

62. Dinesh, D. C. et al. Structural basis of RNA recognition by the SARS-CoV-2 nucleocapsid phosphoprotein. PLOS Pathog. 16, e1009100 (2020).

